# Human Gasdermin D and MLKL disrupt mitochondria, endocytic traffic and TORC1 signaling in budding yeast

**DOI:** 10.1101/2022.11.29.518328

**Authors:** Marta Valenti, María Molina, Víctor J Cid

## Abstract

Gasdermin D (GSDMD) and mixed lineage kinase domain-like protein (MLKL) are the pore-forming effectors of pyroptosis and necroptosis, respectively, with the capacity to disturb plasma membrane selective permeability and induce programmed cell death. The budding yeast *Saccharomyces cerevisiae* has long been used as a simple eukaryotic model for the study of proteins associated with human diseases by heterologous expression. In this work, we expressed in yeast both GSDMD and its N-terminal domain [GSDMD(NT)] to characterize their cellular effects, and compare them to those of MLKL. GSDMD(NT) and MLKL inhibited yeast growth, formed cytoplasmic aggregates, and fragmented mitochondria. Loss-of-function point mutants of GSDMD(NT) showed affinity for this organelle. Besides, GSDMD(NT) and MLKL caused an irreversible cell cycle arrest through TORC1 inhibition, and disrupted endosomal and autophagic vesicular traffic. Our results provide a basis for a humanized yeast platform to study GSDMD and MLKL, a useful tool for structure-function assays and drug discovery.

## Introduction

Pyroptosis and necroptosis are among the programmed cell death mechanisms that guarantee cell survival under circumstances in which internal or external factors compromise tissue or cell homeostasis [1, 2]. The effector of both types of cell death is a pore-forming protein, namely gasdermin D (GSDMD) for pyroptosis and mixed lineage kinase domain-like protein (MLKL) for necroptosis [3–7]. Despite sharing some features, the pathways that lead to their activation and the mechanism by which they permeabilize the plasma membrane differ substantially.

Pyroptosis is elicited upon assembly and activation of nucleotide oligomerization domain (NOD)-like receptors (NLRs) or absent in melanoma-2 (AIM2)-like receptors (AMRs), which are cytosolic innate immune receptors that respond to multiple damage-associated molecular patterns (DAMPs) and pathogen-associated molecular patterns (PAMPs) [8]. These receptors build up a supramolecular organizing center (SMOC) called inflammasome by recruiting the apoptosis-associated speck-like (ASC) adaptor protein and the proinflammatory Caspase-1 protease [9]. GSDMD is constituted by two domains: the N-terminal domain (NTD), which is responsible for the pore-forming activity of the protein and the interaction with membrane lipids through a positively charged region; and the C-terminal domain (CTD), which plays an autoinhibitory role and keeps the protein in an inactive conformation under resting conditions [10, 11]. After inflammasome activation, pro-inflammatory caspases cleave the linker between the NTD and the CTD of GSDMD [3, 4]. Alternatively, Caspase-11 can directly sense cytosolic bacterial lipopolysaccharide and cleave GSDMD [12, 13]. The released NTD endures a conformational change that allows the protein to interact with negatively charged lipids of the plasma membrane, where it forms ring-shaped oligomers and eventually pores [10, 11, 14]. Cells die as a consequence of the loss of membrane selective permeability [7]. However, the plasma membrane is not the only target of GSDMD, as it damages other internal structures, such as mitochondria [15–19] or endosomes [20–22].

On the contrary, different innate immune receptors, including death receptors, Toll-like receptors (TLRs), and receptors for DNA/RNA, trigger necroptosis [23]. In all cases, necroptotic signaling converges in the phosphorylation of the receptor-interacting serine/threonine-protein kinase 3 (RIPK3) [23], which in turn phosphorylates the residues T357/S358 of MLKL [6]. MLKL structure comprises a 4-helix bundle (4HB) domain that is responsible for the interaction with negatively charged lipids of the plasma membrane and oligomerization; a pseudokinase (PK) domain, where the activation loop resides; and a brace that connects these two domains and might also play a role in the interaction with membrane lipids [5, 6, 24, 25]. Phosphorylation of MLKL by RIPK3 induces a conformational change that unleashes the 4HB domain, which can thus interact with plasma membrane lipids, oligomerize and form pores, finally causing cell demise due to the perturbation of cellular homeostasis [5, 26]. Similar to GSDMD, MLKL can also damage other intracellular structures [27–30].

Although the membrane pore-forming activity of GSDMD and MLKL has been extensively studied, some mechanistic details remain poorly characterized, especially regarding their intracellular effects and targets. The budding yeast *Saccharomyces cerevisiae* has been widely used as a simple eukaryotic model to mirror complex aspects of mammalian cell biology [31, 32] due to the high degree of conservation of their molecular pathways and cellular organization [33]. Notably, based on its ready genetic manipulation, researchers have developed a plethora of genetic, genomic, and synthetic biology tools for this model, yielding an alternative platform for the molecular characterization of pathways related to human diseases [34]. Humanized yeast models can be based on the substitution of orthologous genes by their human counterparts [35] or on the integration of human activities or pathways that are naturally lacking in yeast [36, 37]. As a unicellular organism, regulated cell death pathways in yeast are constrained compared to mammalian cells [38], and orthologs to pore-forming effector proteins GSDMD or MLKL are absent. We previously studied human Caspase-1 in the yeast model and demonstrated that it can efficiently recapitulate *in vivo* GSDMD cleavage [39]. Besides, MLKL was recently expressed in budding yeast to establish a model for mechanistic studies on necroptosis [40].

In this work, we aimed to comparatively characterize the performance of the effector proteins of pyroptosis and necroptosis in *S. cerevisiae*. We found that the active form of these proteins inhibits yeast growth, causes cell death, and keeps its capacity to aggregate. However, rather than targeting the plasma membrane, toxicity in yeast is exerted by cell cycle arrest through target-of-rapamycin complex 1 (TORC1) inhibition, alterations of the endosomal traffic and autophagy, and perturbation of the mitochondrial network.

## Results

### The NTD of GSDMD and the 4HB domain of MLKL inhibit yeast growth

In human cells, the NTD of GSDMD, released after Caspase-1-mediated cleavage at D275, is capable of assembling pores through its ability to interact with negatively charged lipids in the plasma membrane, leading to pyroptotic cell death [3, 4]. To establish a yeast model that could be useful to shed some light on open questions in the field, we cloned into *S. cerevisiae* expression vectors cDNAs expressing both the full-length and the NTD (NT) truncated versions of this protein (Fig. 1A) fused to enhanced GFP (EGFP) in C-terminal, both under the control of galactose-inducible *GAL1* promoter. After 5 h of induction in galactose-containing media, we confirmed by immunoblotting that both proteins were expressed (Fig. 1B). However, only GSDMD(NT) strongly impaired yeast growth both in solid (Fig. 1C) and liquid medium, an effect that could be detected early in the exponential growth phase (Fig. 1D). Fusions of GSDMD to a FLAG epitope, as an alternative to the larger EGFP tag, showed the same behavior: GSDMD(NT)-FLAG inhibited yeast growth, whereas full-length GSDMD-FLAG was innocuous (Fig. S1A).

**Figure 1.**
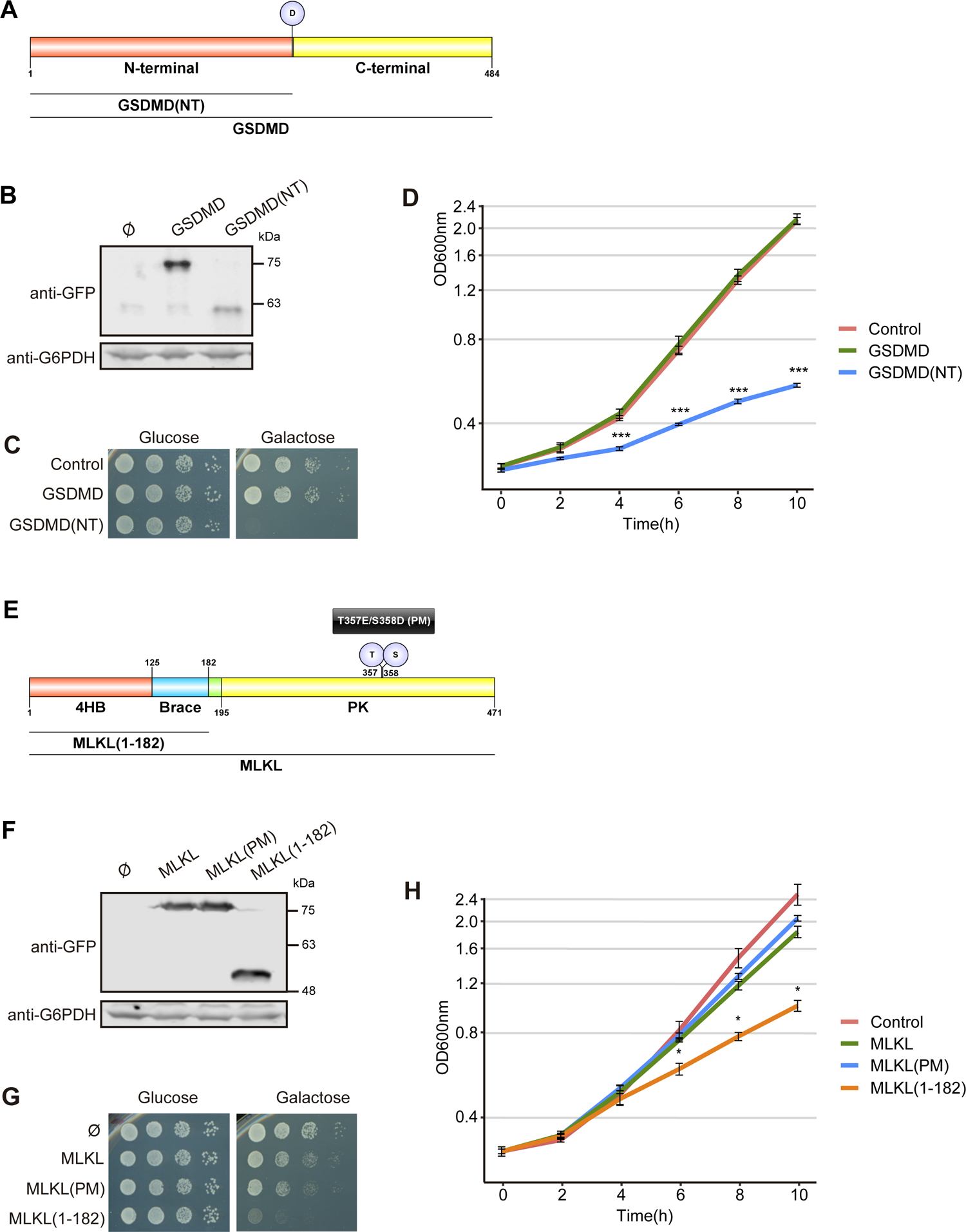
The NTD of GSDMD and the 4HB domain of MLKL inhibit yeast growth. **(A)** Schematic representation of GSDMD depicting its NTD (red), CTD (yellow), and the aspartic residue (D) susceptible to Caspase-1 cleavage. **(B)** Immunoblot showing the expression of GSDMD and GSDMD(NT) in yeast lysates of BY4741 strain bearing plasmids pAG416-GSDMD-EGFP and pAG416-GSDMD(NT)-EGFP after 5 h of induction in SG medium. pAG416-EGFP empty vector was used as a control. The membrane was hybridized with anti-GFP antibody. Anti-G6PDH antibody was used as a loading control. **(C)** Spot growth assay of cells bearing the same plasmids as in (B). Cells were cultured on SD (Glucose) and SG (Galactose) agar media for repression and induction of GSDMD and GSDMD(NT) expression, respectively. **(D)** Growth curves of cells bearing the same plasmids as in (B) performed in SG medium. Measures of optical density at 600nm (OD_600_) were taken each two hours throughout the exponential growth phase. Results are represented as OD_600_ vs. time in a semilogarithmic plot. **(E)** Schematic representation of MLKL depicting its 4HB (red), brace (blue), and PK (yellow) domains. The T357/S358 residues, susceptible to phosphorylation, and their corresponding PMs T357E/S358D are also highlighted. **(F)** Immunoblot showing the expression of MLKL, MLKL(PM) and MLKL(1-182) performed as in (B) using yeast lysates of BY4741 strain bearing plasmids pAG416-MLKL-EGFP, pAG416-MLKL(PM)-EGFP, and pAG416-MLKL(1-182)-EGFP. **(G)** Spot growth assay performed as in (C) but using BY4741 strain bearing the same plasmids as in (F). **(H)** Growth curves of cells bearing the same plasmids as in (F) performed as in (D). A representative assay from three different experiments with different transformants is shown in all cases. In (D, H), results correspond to the mean of three biological replicates performed on different transformants. Error bars represent SD. Asterisks (*, ***) indicate a p-value < 0.05 and 0.001 by Dunn’s test, respectively.

MLKL is a pore-forming protein involved in necroptosis, a different type of programmed cell death in higher cells [5, 7]. Although our primary goal was to model GSDMD activity in yeast, we found it interesting to compare GSDMD(NT) to MLKL, thus delving into putative differences between pyroptosis and necroptosis executioners. MLKL is activated through phosphorylation by RIPK3 in mammalian cells [41, 42]. A recent report showed that human MLKL is not phosphorylated when heterologously expressed in yeast unless it is co-expressed with RIPK3 [40]. To by-pass the phosphorylation step, we cloned both the wild-type *MLKL* gene and a phosphomimetic T357E/S358D version, referred hereafter to as MLKL(PM), in the same expression vector used for GSDMD (Fig. 1E). Both proteins were efficiently produced in yeast (Fig. 1F), reaching much higher levels of expression than GSDMD (Fig. S1B), and their expression led to mild growth inhibition (Fig. 1G-H). Some controversy has arisen about whether MLKL phosphorylation is sufficient to activate this protein, with studies using the phosphomimetic mutant protein yielding contradictory results [27, 41, 43]. Our results in yeast show no significant differences in growth inhibition between phosphomimetic (PM) and wild-type (WT) versions of MLKL. Previous reports state that the 4HB domain located at the N-terminus of this protein is responsible and sufficient for the interaction of MLKL with membrane lipids and subsequent permeabilization of the membrane, while the PK domain might play a regulatory and/or autoinhibitory role [5, 44]. To test this, we cloned the NTD of MLKL, comprising the 4HB plus the brace region, hereafter referred to as MLKL(1-182), in the same vector used for the other MLKL and GSDMD constructs (Fig. 1E). The level of expression of this truncated form of MLKL was comparable to those of the full-length WT and PM versions (Figs. 1F and S1B), but it exerted higher growth inhibition, as reflected in Fig. 1G-H.

Overall, these results prove that both GSDMD and MLKL are functional in our experimental setting, allowing us to establish an *in vivo* model to explore their activity and mechanism of action. Interestingly, the effect of GSDMD(NT) on yeast growth is more severe *per se* than that of the MLKL 4HB domain. Also, our model recapitulates the autoinhibitory function of the C-terminal extensions of both proteins, which is tighter in the case of GSDMD as compared to MLKL.

### In yeast, the NTD of GSDMD and MLKL aggregate in cytoplasmic spots and reduce cell viability, but do not cause severe cell lysis

Once activated, GSDMD and MLKL are known to insert into cellular membranes through positively charged patches on their surface that allow them to interact with negatively charged lipids, particularly cardiolipin, phosphatidylserine, and phosphoinositides [4–6, 10]. Previously, we reproduced recognition of the plasma membrane by positively charged human proteins involved in innate immune signaling in yeast, like Toll/interleukin-1 receptor domain-containing adapter protein (TIRAP) [45]. Thus, we expected yeast growth inhibition by GSDMD and MLKL to be linked to their localization at the plasma membrane, leading to its disruption. However, none of the GSDMD and MLKL EGFP fusions produced in yeast was detected at the plasma membrane by fluorescence microscopy (Fig. 2A-B). Full-length GSDMD showed a diffuse nucleo-cytoplasmic pattern typical of soluble proteins, while GSDMD(NT) formed small foci distributed throughout the cell cytoplasm. In the case of MLKL, we observed that the WT and PM constructs formed one or two larger, brighter spots per cell. However, in MLKL(1-182)-expressing cells, bright spots were less frequent and were substituted by small numerous foci, similar to the ones observed for GSDMD(NT). Immunoblotting on lysates from these cultures processed under non-reducing conditions revealed an enhanced presence of high molecular weight protein aggregates in cells bearing GSDMD(NT) and the different constructs of MLKL, but not in the case of full-length GSDMD (Fig. 2C-D).

**Figure 2.**
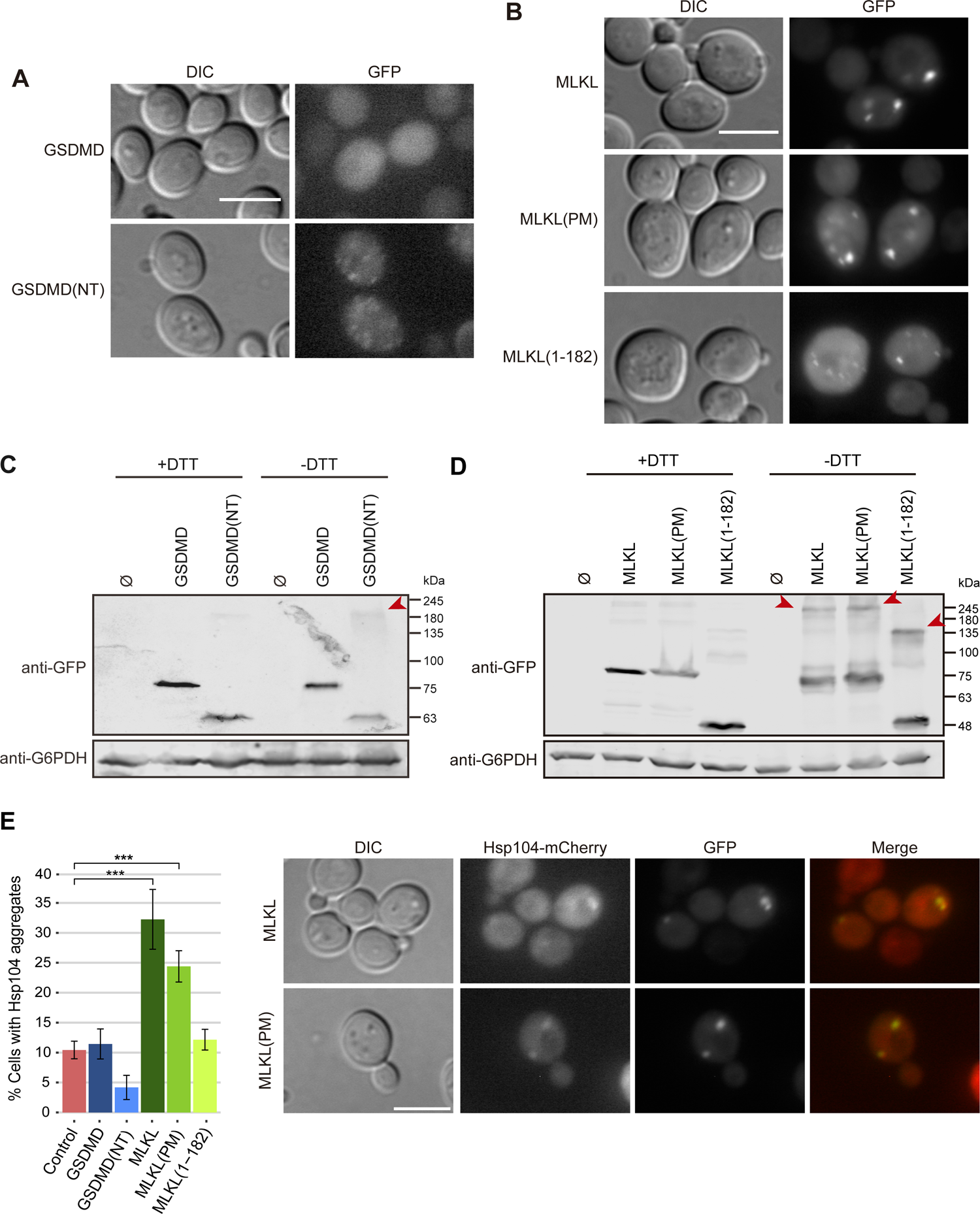
The NTD of GSDMD and MLKL aggregate in cytoplasmic spots in yeast. **(A-B)** Fluorescent and bright-field differential interference contrast (DIC) microscopy of BY4741 strain bearing the same plasmids as in Fig. 1B and Fig. 1F, respectively. **(C-D)** Immunoblots showing a comparison under reducing (+DTT) and non-reducing (-DTT) conditions of yeast lysates of BY4741 strain bearing the same plasmids as in A-B, respectively. Red arrowheads mark high molecular weight protein aggregates enhanced in the absence of DTT. The membranes were hybridized with anti-GFP antibody. Anti-G6PDH antibody was used as a loading control. **(E, left panel)** Graph showing the percentage of cells with Hsp104 aggregates (n>100) for each population of MVY07 strain bearing the same plasmids as in A-B. Results correspond to the mean of three biological replicates performed on different transformants. Error bars represent SD. Asterisks (***) indicate a p-value < 0.001 by Tukey’s HSD test. Only statistical differences between the different samples and the control are depicted. **(E, right panel)** Colocalization of Hsp104 aggregates with MLKL and MLKL(PM), respectively. Protein expression was induced for 5 h in SG medium in all cases. All scale bars indicate 5 µm. A representative assay from three different experiments with different transformants is shown in all cases.

The observed cytoplasmic large bright spots formed by MLKL-EGFP and MLKL(PM)-EGFP might reflect the accumulation of misfolded protein aggregates within the cell. To assess this possibility, we transformed the different GSDMD and MLKL-producing plasmids in a yeast strain in which Hsp104, one of the main chaperones involved in the formation of different protein bodies [46], was tagged with the fluorescent protein mCherry. Under basal conditions, Hsp104 remains soluble in the nucleus and cytoplasm of the cells. When cells are subdued to a change in cellular homeostasis, this protein relocates to proteostatic stress compartments [46]. Both MLKL and MLKL(PM) induced the formation of Hsp104 aggregates, while GSDMD, GSDMD(NT), and MLKL(1-182) did not (Fig. 2E, left panel). In the case of MLKL, 74±11% of Hsp104 aggregates colocalized with MLKL spots; and in the case of MLKL(PM), 69±12% of Hsp104 aggregates colocalized with MLKL(PM) spots (Fig. 2E, right panel). Thus, MLKL induces the formation of proteostatic stress compartments when overexpressed in yeast, whereas the more toxic GSDMD(NT) and MLKL(1-182) neither trigger proteostatic stress nor co-localize with Hsp104.

If pore formation in the plasma membrane was the cause of strong growth inhibition of GSDMD(NT) and MLKL(1-182) in yeast, we should expect severe cell lysis to occur. Even though neither GSDMD(NT)-EGFP nor MLKL(1-182)-EGFP seemed to associate with the yeast plasma membrane as observed by fluorescence microscopy, we performed propidium iodide (PI) staining as a readout of putative loss of plasma membrane selective permeability and analyzed the cultures by flow cytometry (Fig. 3A-B). Although there was a significant increase in the percentage of PI-positive cells both for GSDMD(NT) and MLKL(1-182) compared to the negative control or their full-length counterparts, the overall percentage of lysed cells was too low (<8%) after 5h of induction. At longer incubation times (12 h post-induction) this percentage increased 3 to 5-fold for all transformant cells, particularly in the case of GSDMD(NT) (Fig. S2A).

**Figure 3.**
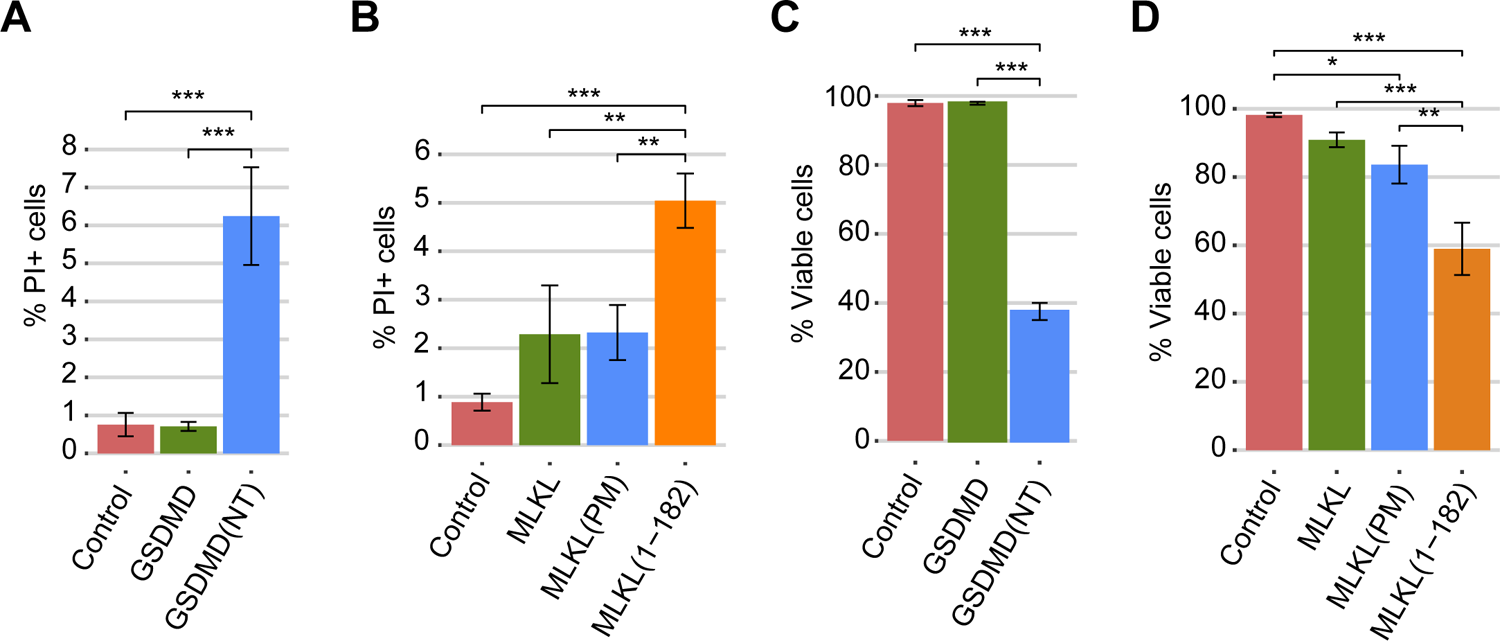
The NTDs of GSDMD and MLKL cause a decrease in cell viability. **(A-B)** Graphs showing the percentage of PI-positive stained cells (n=10,000) for each population of BY4741 strain bearing the same plasmids as in Fig. 1B and Fig. 1F, respectively. **(C-D)** Graphs showing the percentage of viable cells determined by a cell viability assay of BY4741 strain bearing the same plasmids as in (A-B), respectively. Protein expression was induced for 5 h in SG medium in all cases. Results correspond to the mean of three biological replicates performed on different transformants. Error bars represent SD. Asterisks (*, **, ***) indicate a p-value <0.05, <0.01, and <0.001, respectively, by Tukey’s HSD test.

A previous report highlighted that the fusion of a bulky C-terminal tag in GSDMD(NT) might reduce the efficiency of pyroptosis [47]. To determine whether the EGFP tag might be hindering GSDMD(NT) interaction with the plasma membrane, we performed the same experiment with a C-terminal FLAG fusion. Indeed, the percentage of PI-positive GSDMD(NT)-FLAG-expressing cells was significantly higher as compared to that of cells producing GFP-tagged GSDMD(NT) (Fig. S2B), reaching 25% of the population after 5h of induction. Thus, the EGFP tag is likely preventing cell lysis by interfering with GSDMD(NT) translocation to the plasma membrane in yeast.

Given that the severity of growth inhibition induced by GSDMD(NT)-EGFP and MLKL(1-182)-EGFP at early time points (Fig. 1D and H) did not correlate with the mild increase in the percentage of lysed PI-positive cells (Fig. 3A-B), we were prompted to examine cell viability through a microcolony formation assay. We observed a decrease in viability after 5 h of induction in galactose-containing media, particularly significant in the case of GSDMD(NT) and MLKL(1-182) (Fig. 3C-D), which follows the same trend as growth inhibition. Altogether, these results indicate that GSDMD(NT)-EGFP and MLKL(1-182)-EGFP cause a reduction of cell viability soon after induction by a mechanism that differs from the permeabilization of the plasma membrane mechanism expected for pore-forming effectors. But if GSDMD(NT)-EGFP is not causing cell lysis, why is it so toxic for yeast cells? We hypothesized that accumulation of GSDMD(NT)-EGFP in intracellular membranes was responsible for the damage leading to severe loss of viability, making this setting suitable for the study of the effects of GSDMD on its cytoplasmic targets.

### The NTDs of GSDMD and MLKL alter the yeast mitochondrial network

Different reports prove that GSDMD(NT) can interact with mitochondria and cause mitochondrial depolarization, fragmentation, and release of mitochondrial DNA to the cytosol by a yet undefined mechanism [15–17, 19]. MLKL causes a similar effect on mitochondria [6, 30]. We questioned whether the mitochondrial network was affected in the yeast model. For this purpose, we co-expressed the two GSDMD versions with the mitochondrial marker Ilv6-mCherry and visualized cells by confocal fluorescence microscopy. As in mammalian cell lines, around 40% of yeast cells expressing GSDMD(NT)-EGFP showed fragmented mitochondria, but the majority of GSDMD(NT)-EGFP spots did not colocalize with them (Fig 4A-B and Fig. S3A). However, we could not detect significant changes in mitochondrial membrane potential or ROS levels (data not shown).

**Figure 4.**
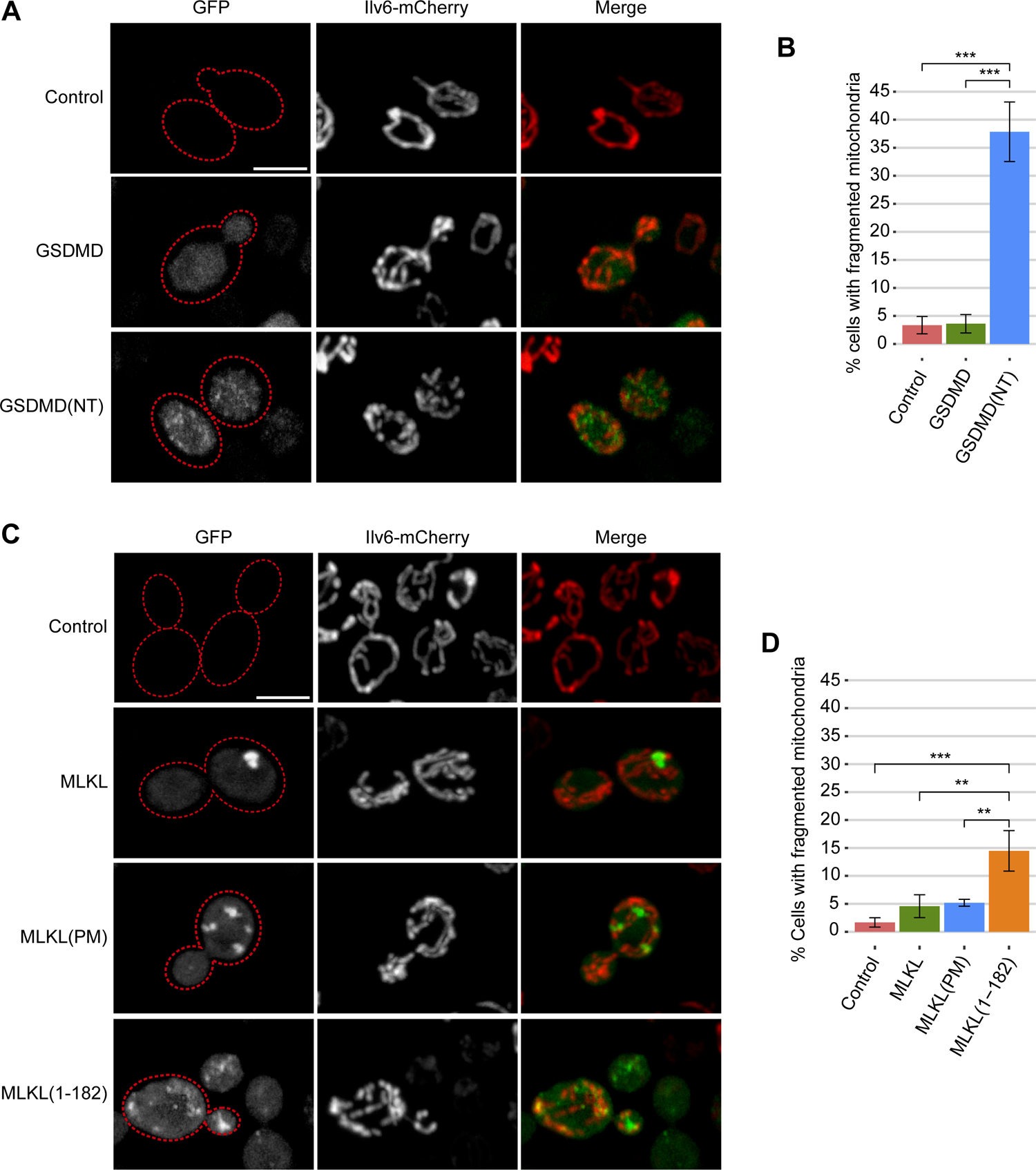
The NTDs of GSDMD and MLKL fragment the mitochondrial network. **(A)** Stacked images obtained by confocal fluorescence microscopy of BY4741 strain bearing the mitochondrial marker pOB06 (Ilv6-mCherry) and plasmids pAG416-GSDMD-EGFP and pAG416-GSDMD(NT)-EGFP, respectively. pAG416 empty vector was used as a control. See also Fig. S3A. **(B)** Quantification (n>100) by fluorescence microscopy of the percentage of cells showing fragmented mitochondria for each population shown in (A). **(C)** Stacked images obtained by confocal fluorescence microscopy of BY4741 strain bearing the mitochondrial marker pOB06 (Ilv6-mCherry) and plasmids pAG416-MLKL-EGFP, pAG416-MLKL(PM)-EGFP, and pAG416-MLKL(1-182)-EGFP, respectively. pAG416 empty vector was used as a control. See also Fig. S3B. **(D)** Quantification (n>100) by fluorescence microscopy of the percentage of cells showing fragmented mitochondria for each population shown in (C). Protein expression was induced for 5 h in SG medium. All scale bars indicate 5 µm. Results correspond to the mean of three biological replicates performed on different transformants in all cases. Error bars represent SD. Asterisks (*, **, ***) indicate a p-value <0.05, <0.01, and <0.001 by Tukey’s HSD test.

Then, we made the same confocal microscopy experiments with the different constructs of MLKL. The effect of MLKL(1-182) on the mitochondrial network was less severe than that of GSDMD(NT), as only 15% of the cells had disrupted mitochondria (Fig. 4C-D and Fig. S3B). However, contrary to GSDMD(NT), MLKL(1-182) spots frequently colocalized with mitochondria. As for full-length MLKL and MLKL(PM), they neither impacted the mitochondrial network nor colocalized with them (Fig. 4C-D and Fig. S3B). Thus, as reported in mammalian cells, the yeast mitochondrial network is targeted by the NTDs of GSDMD and MLKL, providing a plain model to study their interaction with this organelle.

### Key point mutations at interaction interfaces of the NTD of GSDMD abrogate cytoplasmic aggregates and growth inhibition

After cleavage, GSDMD(NT) monomers undergo a conformational change that allows them to bind membrane lipids and oligomerize. Liu *et al.* [11] described that this oligomerization process is driven through three interaction interfaces and found critical residues within those interfaces for pyroptotic activity in mice. To challenge our yeast model for functional studies on the human protein and, particularly, to evaluate the intracellular consequences of GSDMD mutation, we selected one mutation for each interface (L60G for interface I; F81D for interface II; and I91D for interface III) and mutated the equivalent amino acids in human GSDMD(NT)-EGFP (L59G, F80D, and I90D, respectively). See Fig. S4A-B for human-mouse GSDMD(NT) alignment and location of the residues in the tertiary structure of homologous mouse GSDMA3. Liu *et al.* [11] also described the interactions between the NTD and CTD of mouse GSDMD that maintain the protein inactive under basal conditions and characterized mutants that acquired spontaneous pyroptotic activity due to alterations in such interactions [11]. We selected among them the one that most enhanced pyroptotic markers, A380D, and mutated the equivalent residue in human full-length GSDMD-EGFP (A377D). Finally, GSDMD(NT) interacts with plasma membrane lipids through a positively charged patch on its surface formed by four basic residues (R138/K146/R152/R154 in mouse GSDMD). Replacement of those four residues by alanine blocks pyroptosis because it hinders the assembly of the pores [10]. To explore the effect of these mutations in our model we produced the equivalent human GSDMD(NT)-EGFP quadruple mutant (R137A/K145A/R151A/R153A), hereafter referred to as 4A. As shown in Fig. 5A, mutation of residues identified as part of interaction interfaces I (L59G) and II (F80D) of GSDMD(NT)-EGFP monomers, as well as the blockade of interaction with membrane phospholipids (4A), were no longer inhibitory for yeast growth. On the contrary, the mutation of I90, which belongs to interface III, did not alter its toxicity. These mutants showed a similar behavior when the EGFP tag was replaced by a FLAG epitope, with the only difference that the I90D mutant displayed partial loss-of-function (Fig. S4C), even though it failed to permeabilize the plasma membrane after 5 h of induction, measured by PI uptake (Fig. S4D). These results suggest that human GSDMD(NT) monomers recapitulate their interactions among them and with lipid surfaces in yeast, and that such mechanisms are conserved between the human protein and its mouse homolog, at least for interfaces I and II. Interface III might play a secondary role, might be less critical for the formation of polymers that interfere with yeast essential cellular functions, or may not be as crucial for human GSDMD(NT) as for the murine protein. Finally, full-length GSDMD A377D did not gain spontaneous activity in yeast cells since it behaved like WT GSDMD in growth assays (Fig. 5A). All the mutant versions seemed stable in yeast and were expressed in similar levels, as determined by immunoblotting (Fig. 5B).

**Figure 5.**
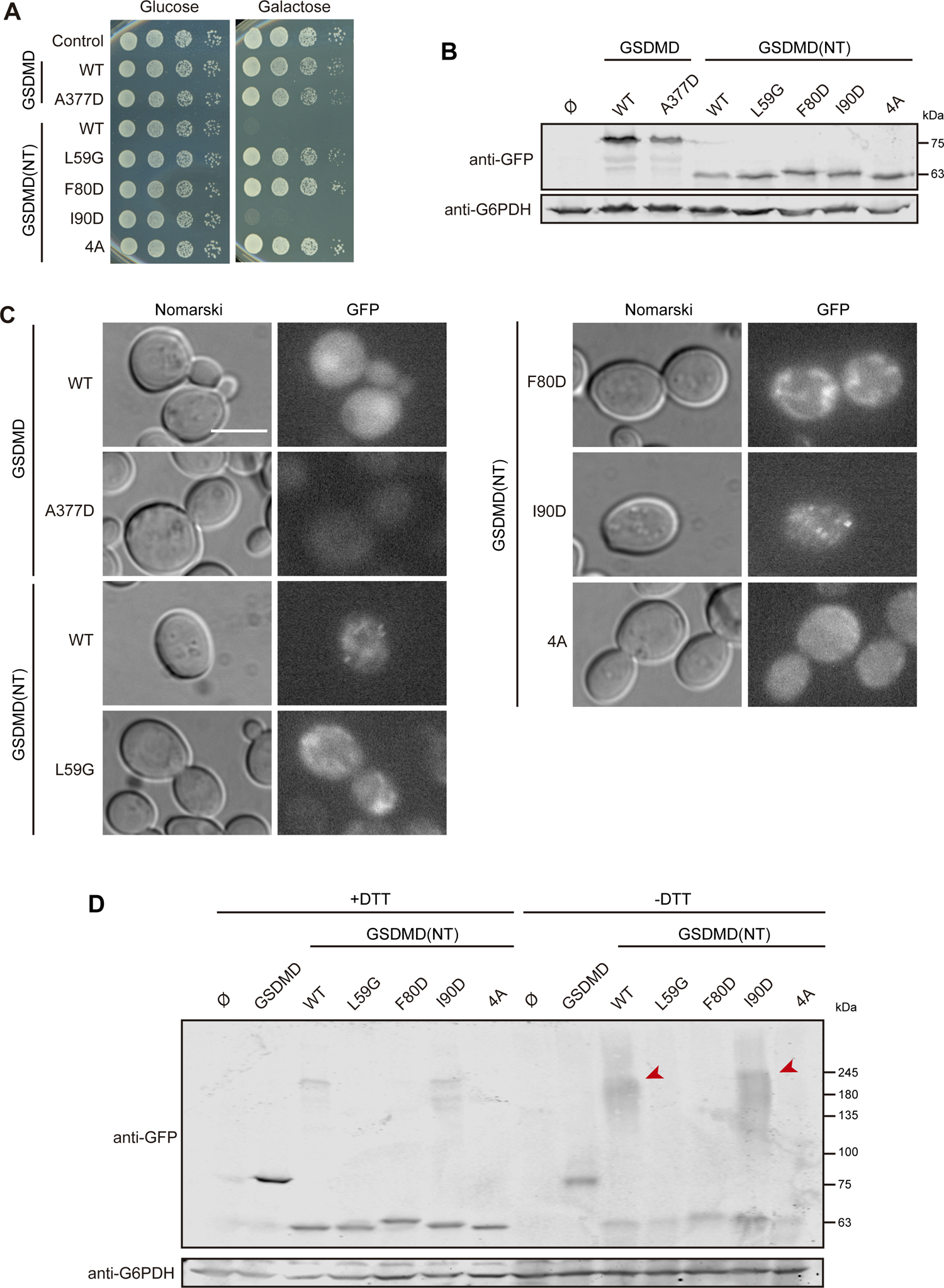
Functionality correlates with aggregation in mutants of the NTD of GSDMD. **(A)** Spot growth assay of BY4741 strain bearing plasmids pAG416-GSDMD-EGFP WT or A377D and pAG416-GSDMD(NT)-EGFP WT, L59G, F80D, I90D, or 4A. Cells were cultured on SD (Glucose) and SG (Galactose) agar media for repression and induction of GSDMD and GSDMD(NT) specified versions, respectively. pAG416-EGFP empty vector was used as a control. **(B)** Immunoblot showing the expression of GSDMD and GSDMD(NT) mutants in yeast lysates of BY4741 strain bearing the same plasmids as in (A) after 5 h of induction in SG medium. **(C)** Fluorescent and bright field (DIC) microscopy of BY4741 strain bearing the same plasmids as in (A) after 5 h of induction in SG medium. Scale bar indicates 5 µm. **(D)** Immunoblot showing a comparison under reducing (+DTT) and non-reducing (-DTT) conditions of yeast lysates of BY4741 strain bearing the same plasmids as in A, after 5 h of induction in SG medium. Red arrowheads mark high molecular weight protein aggregates enhanced in the absence of DTT. Membranes in (B, D) were hybridized with anti-GFP antibody. Anti-G6PDH antibody was used as a loading control. A representative assay from three different experiments with different transformants is shown in all cases.

Next, we evaluated possible changes in protein localization and aggregation of the different mutant proteins by fluorescence microscopy and immunoblotting in non-reducing conditions, respectively. As shown in Fig. 5C, the subcellular distribution of non-functional mutants (L59G, F80D, and 4A) was neither in small numerous foci like the WT GSDMD(NT) nor uniformly nucleocytoplasmic like full-length GSDMD. Rather, they seemed to be diffusely attached to intracellular structures, although in the case of GSDMD(NT) 4A, this pattern was less pronounced. By contrast, the functional mutant I90D did not show differences compared to GSDMD(NT) WT, forming numerous small spots within the cells. As for full-length GSDMD A377D, it showed a diffuse distribution. Finally, immunoblotting in non-reducing conditions revealed, as expected, that only functional proteins [i.e., GSDMD(NT) WT and I90D] formed higher-order oligomers (Fig. 5D). The behavior of these different loss-of-function mutants further underscore that there is a strong correlation between growth inhibition and aggregation of GSDMD(NT) in yeast.

### Non-functional mutants of the NTD of GSDMD colocalize with the mitochondrial network

We have shown above that GSDMD(NT) interferes with yeast mitochondria, so we aimed to verify that loss-of-function GSDMD(NT) mutants failed to disturb this organelle. For this purpose, we co-expressed the corresponding mutants with the mitochondrial marker Ilv6-mCherry and visualized the cells by confocal microscopy. As predicted, the mitochondrial network of cells expressing GSDMD(NT) L50G, F80D and 4A was intact, while that of cells expressing GSDMD(NT) I90D was fragmented (Fig. 6 and Fig. S5). Interestingly, the EGFP fusions of non-functional mutants of GSDMD(NT), namely L50G, F80D, and, to a lesser extent, 4A, colocalized with the mitochondrial network under basal conditions. These data suggest that, when GSDMD(NT) fails to homopolymerize, the individual monomers associate to mitochondrial membranes.

**Figure 6.**
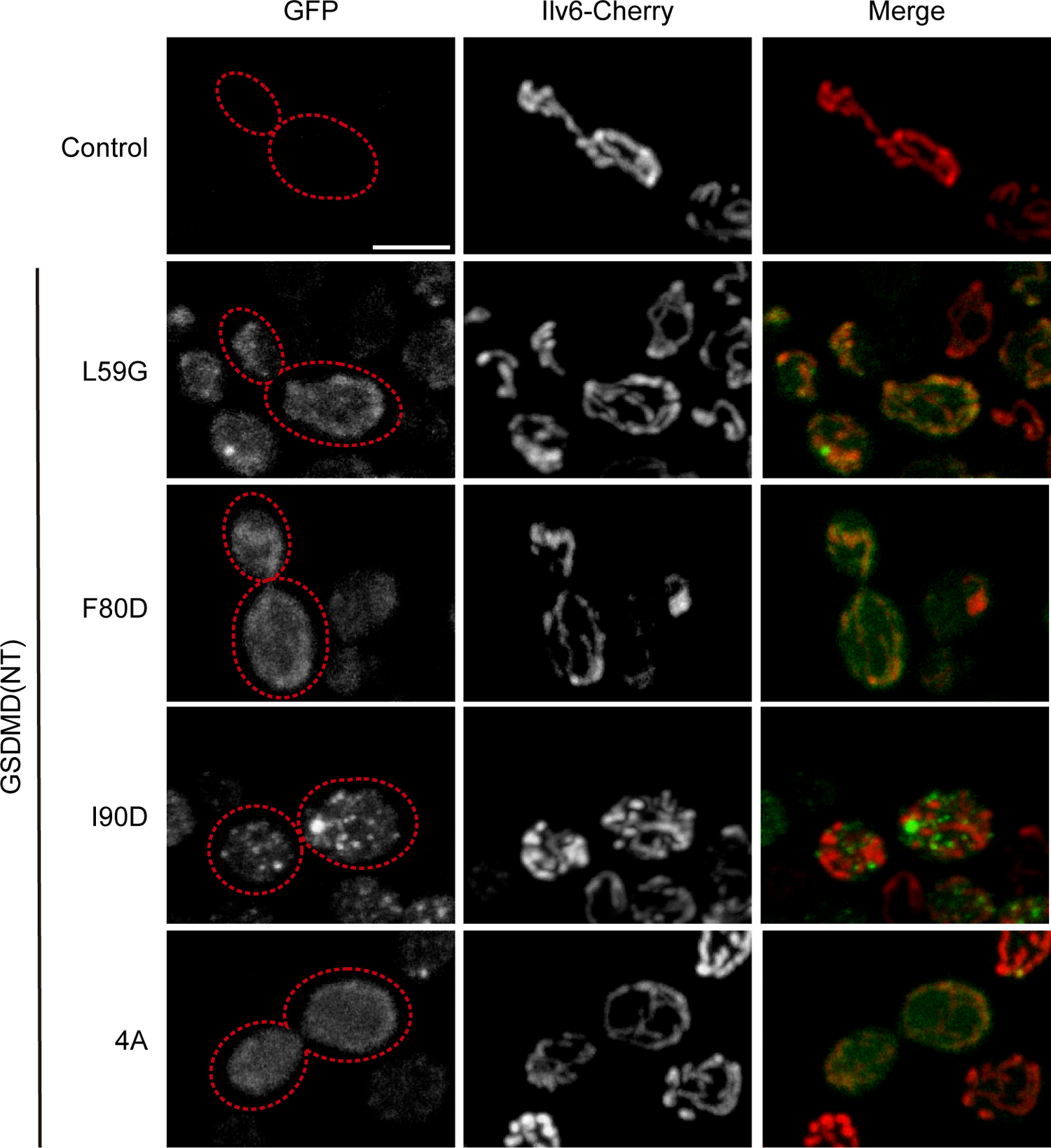
Non-functional mutants of the NTD of GSDMD colocalize with the mitochondrial network. Stacked images obtained by confocal fluorescence microscopy of BY4741 strain bearing the mitochondrial marker pOB06 (Ilv6-mCherry) and plasmids pAG416-GSDMD(NT)-EGFP L59G, F80D, I90D or 4A, after 5 h of induction in SG medium. pAG416 empty vector was used as a control. Scale bar indicates 5 µm. See also Fig. S5.

### The NTDs of GSDMD and MLKL cause cell cycle arrest through inhibition of TORC1

Although the effects of the NTDs of GSDMD and MLKL on yeast mitochondria are significant, they are not severe enough to account for the strong growth inhibition and loss of viability observed, especially in the case of GSDMD(NT)-expressing cells. We hypothesized that additional factors should be contributing to the resulting phenotype. In the previous microscopy experiments, we noticed the presence of an unusual fraction of unbudded cells, especially when we expressed GSDMD(NT). To address whether the growth defect observed was linked to an arrest in cell cycle progression, we induced the expression of all GSDMD and MLKL versions for 5 h in galactose-containing media, and then analyzed cellular DNA content by flow cytometry. As shown in Fig. 7A, GSDMD(NT) and MLKL(1-182) caused a statistically significant increase in the percentage of cells in G1 phase (non-replicated DNA content) in asynchronous cultures compared to control cells or to their respective full-length versions. Besides, this percentage was higher for GSDMD(NT) than for MLKL(1-182), in correlation with their respective growth inhibition and effect on cell viability. These results were obtained by expressing fusions to EGFP, but GSDMD(NT) fused to FLAG induced a similar phenotype (Fig. S6A), dismissing the possibility of an artifact caused by the epitope. To confirm a suspected G0/G1 cell cycle arrest, we stained yeast cells expressing GSDMD and GSDMD(NT) with rhodamine-conjugated phalloidin (Rd-phalloidin) to visualize the actin cytoskeleton, which supports polarized growth for budding. As expected for a cell cycle arrest in G0/G1, we observed an increase in the percentage of unbudded cells as well as a decrease in the percentage of cells with a polarized cytoskeleton among the unbudded cells (i.e., cells ready to start a new round of cell cycle) when GSDMD(NT) was expressed, as compared to control or GSDMD-expressing cells (Fig. S6B), indicating an arrest in cell cycle progression. MLKL(1-182) induced a similar effect (Fig. S6C).

**Figure 7.**
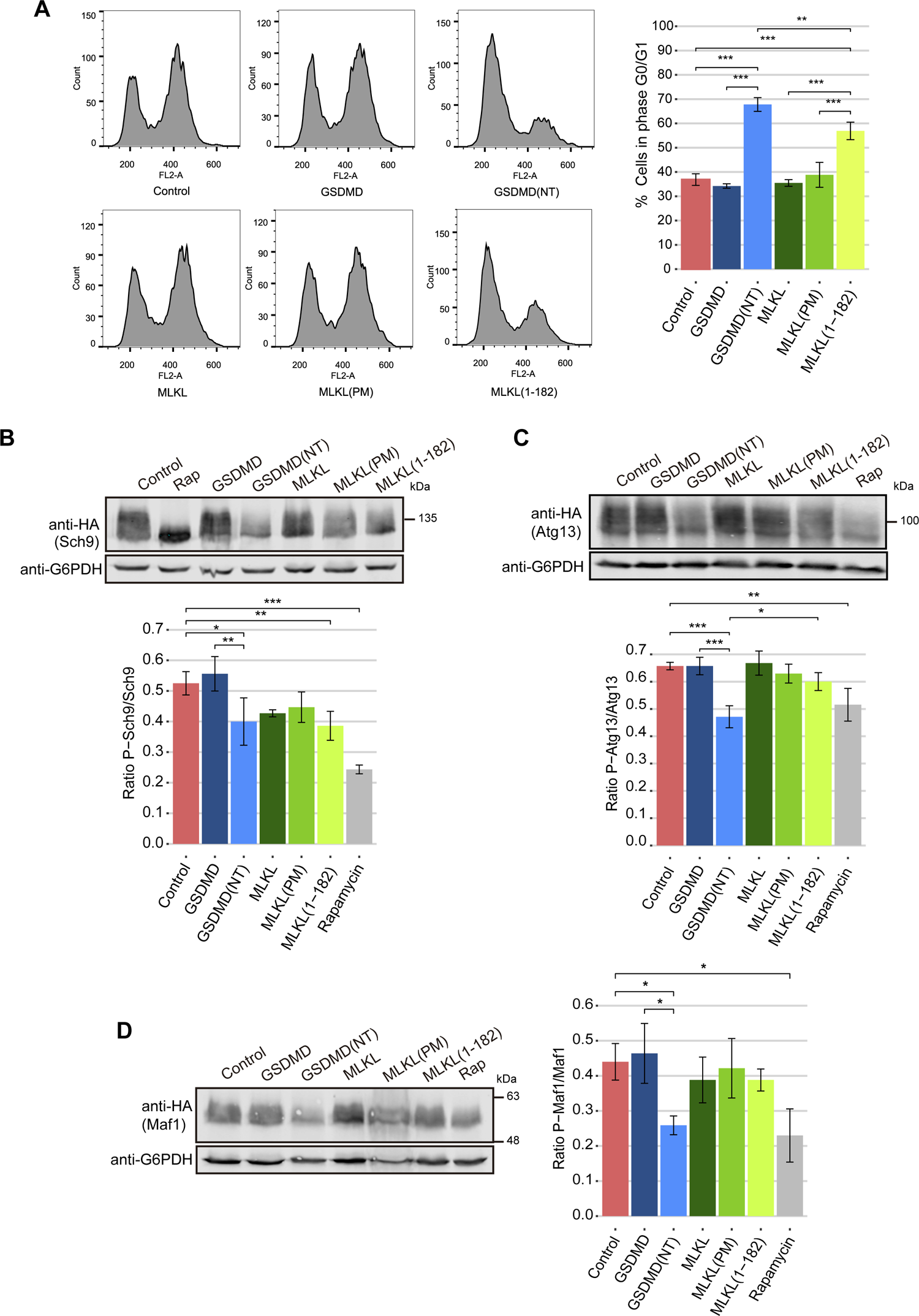
The NTDs of GSDMD and MLKL cause cell cycle arrest through the inhibition of TORC1. **(A)** Cell cycle profiles obtained by measuring DNA content (FL2-A) of cells stained with PI and subsequently analyzed by flow cytometry (n=10,000) (left panels), and graph showing the percentage of cells in phase G0/G1 for each population (right panel) of BY4741 strain bearing plasmids as in Fig. 1B and Fig. 1F, after 5 h of induction in SG medium. **(B)** Immunoblot showing Sch9 phosphorylation (upper panel) and quantification of P-Sch9 relative to total Sch9 (lower panel) in yeast lysates of BY4741 strain bearing the plasmid pJU676 (Sch9-5xHA) and pAG413-GSDMD-EGFP, pAG413-GSDMD(NT)-EGFP, pAG413-MLKL-EGFP, pAG413-MLKL(PM)-EGFP or pAG413-MLKL(1-182)-EGFP after 7 h of induction in SG medium. pAG413-EGFP empty vector was used as a control. **(C)** Immunoblot showing Atg13 phosphorylation (upper panel) and quantification of P-Atg13 relative to total Atg13 (lower panel) in yeast lysates of BY4741 strain bearing the plasmid HC078 (3xHA-Atg13) and the same plasmids as in (B). **(D)** Immunoblot showing Maf1 phosphorylation (upper panel) and quantification of P-Maf1 relative to total Maf1 (lower panel) in yeast lysates of BY4741 strain bearing the plasmid pAH099 (Maf1-3xHA) and the same plasmids as in (B). In (B-D), cells treated with 100nM rapamycin for 5 h were used as a positive control of TORC1 inhibition. Membranes were hybridized with anti-HA antibody. Anti-G6PDH antibody was used as a loading control. A representative blot from three different experiments with different transformants is shown. In (A-D), results correspond to the mean of three biological replicates performed on different transformants. Error bars represent SD. Asterisks (*, **, ***) indicate a p-value<0.05, <0.01, and <0.001, respectively, by Tukey’s HSD test.

The TORC1 complex, homolog to mammalian mTORC1, is one of the core regulators of cell cycle and growth in yeast [48]. Yeast TORC1 senses the concentration of amino acids available in the medium and regulates yeast growth accordingly. In the presence of nutrients, the regulatory exit from G0 complex (EGOC) interacts and activates TORC1, and the kinases of this complex, Tor1/Tor2, phosphorylate their substrates to promote cell proliferation. Under starvation conditions, TORC1 is inhibited and halts cell growth [49]. Previous works have used the electrophoretic mobility shift caused by phosphorylation of Sch9, one of the main targets of TORC1, as a readout to evaluate TORC1 activity [50]. This protein controls ribosome biogenesis, protein translation, and cell cycle progression [50]. To determine whether TORC1 inhibition was the mechanism underlying cell cycle arrest in yeast cells expressing GSDMD(NT) and MLKL(1-182), we co-expressed the plasmids carrying these constructs with a plasmid expressing an Sch9-HA fusion. Cells transformed with an empty vector were used as a negative control of TORC1 inhibition, while cells treated with rapamycin were used as a positive control. As shown in Fig. 7B, the Sch9 mobility shift observed in control cells, completely disappeared in the presence of rapamycin. A significant decrease in Sch9 phosphorylation was also observed in cells expressing either GSDMD(NT) or MLKL(1-182) compared to control cells, indicative of TORC1 inhibition.

TORC1 inhibition induces several adaptations for survival under nutrient depletion in the yeast cell, including the inhibition of transcription [51] and the induction of autophagy [52]. To corroborate our results, we evaluated possible changes in RNA transcription by measuring the phosphorylation of Maf1, a negative regulator of RNA polymerase III that is phosphorylated by TORC1 [53, 54]. Similarly, we assessed changes in autophagy signaling by measuring Atg13 phosphorylation. TORC1 inhibition leads to its dephosphorylation, triggering autophagy [52]. Equivalently to Fig 7B, we co-transformed the plasmids carrying GSDMD and MLKL constructs with plasmids expressing either Maf1-HA or HA-Atg13. Cells bearing an empty vector were used as a negative control, while cells treated with rapamycin were used as a positive control. Surprisingly, while Maf1 and Atg13 did become dephosphorylated in cells expressing GSDMD(NT) as compared to control cells, there was no significant effect in cells expressing MLKL(1-182) (Fig. 7C-D). This could mean that the mechanism by which these proteins interfere with TORC1 signaling differs. Altogether, our results show that both GSDMD(NT) and MLKL(1-182) cause a cell cycle arrest through TORC1-Sch9 signaling pathway inhibition, but differ in the effect on RNA transcription and autophagic signaling. This could explain the differences in the magnitude of the cell growth defect induced by GSDMD(NT) and MLKL(1-182).

### GSDMD(NT) and MLKL disrupt autophagic flux

As stated above, the decrease of TORC1-imposed Atg13 phosphorylation is the signal that triggers the autophagic flux [52]. We decided to test by immunoblotting whether GSDMD(NT) effectively induced this pathway, using Atg8-GFP degradation as a marker [55]. When autophagy is functional, Atg8-GFP is transported together with the autophagosome to the vacuoles (equivalent to mammalian lysosomes) and degraded, releasing a GFP moiety that can be visualized as a ≈25 kDa band with anti-GFP antibodies in immunoblots. Contrary to what we expected, expression of GSDMD(NT) failed to significantly induce autophagy as compared to a rapamycin-treated control (Fig. 8A). Since autophagy is crucial for cell survival under nitrogen starvation conditions, and the blockade of this pathway when TORC1 is inhibited leads to loss of viability [56, 57], this result might explain the loss of viability observed in Fig. 3C for GSDMD(NT)-expressing cells.

**Figure 8.**
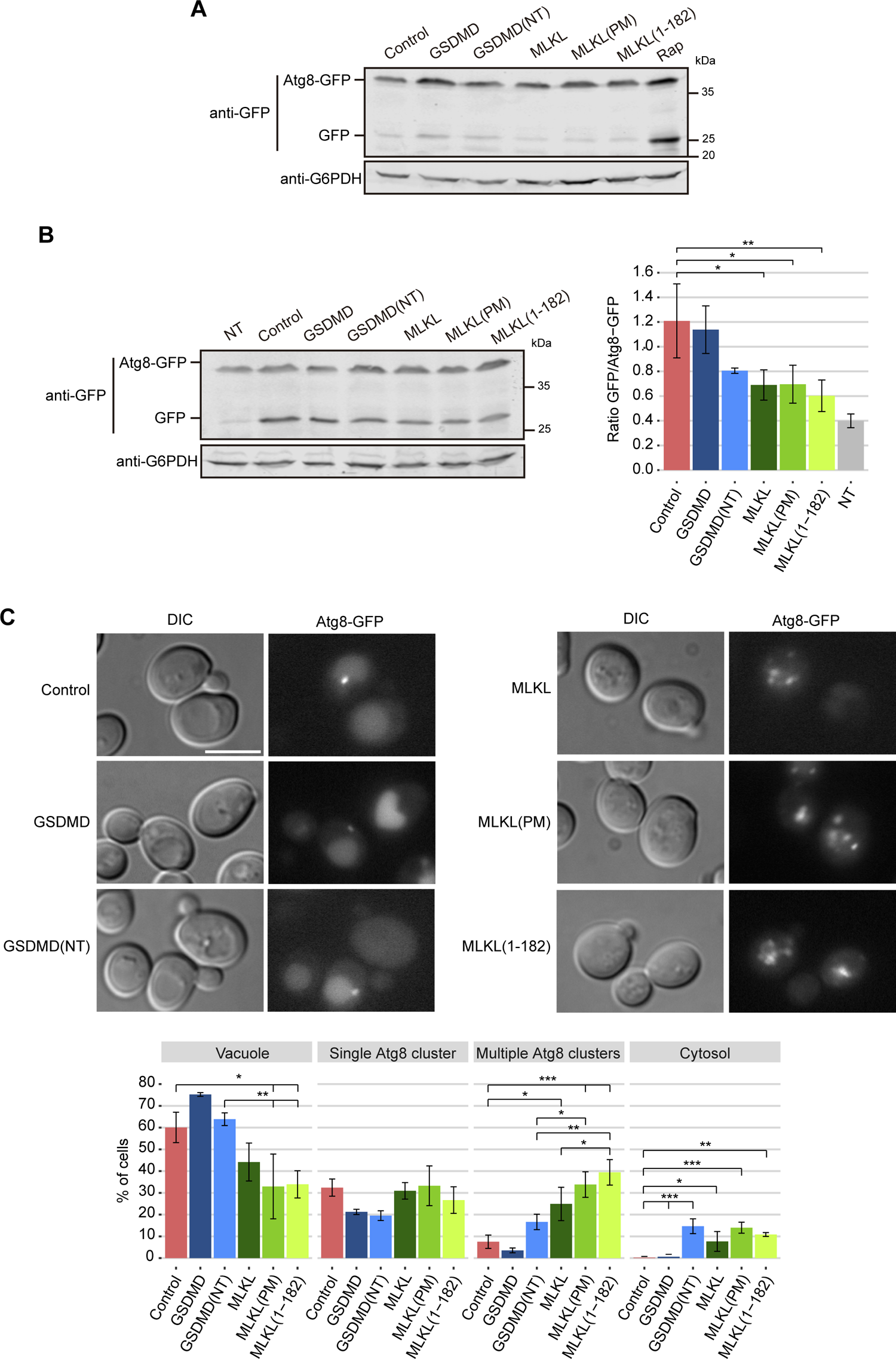
GSDMD(NT) and MLKL impair autophagy. **(A)** Immunoblot showing Atg8-GFP degradation in yeast lysates of BY4741 *trp1Δ* strain bearing the autophagic marker pRS314-GFP-Atg8 and pAG413-GSDMD-DsRed, pAG413-GSDMD(NT)-DsRed, pAG413-MLKL-DsRed, pAG413-MLKL(PM)-DsRed or pAG413-MLKL(1-182)-DsRed after 7 h of induction in SG medium. pAG413-DsRed empty vector was used as a control. Cells treated with 100nM rapamycin for 5 h were used as a positive control of autophagy. **(B)** Immunoblot showing Atg8-GFP degradation (left panel) and quantification of released GFP relative to Atg8-GFP (right panel) after 5 h of induction in SG medium followed by a 2 h treatment with rapamycin 100nM in yeast lysates of BY4741 *trp1Δ* strain bearing the same plasmids as in (A). Cells bearing pAG413-DsRed empty vector and treated with rapamycin for 2 h were used as a positive control of autophagy. Cells bearing pAG413-DsRed empty vector and untreated with rapamycin (NT) were used as a negative control of autophagy. **(C)** Fluorescent and bright-field (DIC) (upper panels) and quantification (n>100) of Atg8-GFP localization (lower panel) performed as in (B). Scale bar indicates 5 µm. In (A-B), membranes were hybridized with anti-GFP antibody. Anti-G6PDH antibody was used as a loading control. A representative blot from three different experiments with different transformants is shown. In (B-C), results correspond to the mean of three biological replicates performed on different transformants. Error bars represent SD. Asterisks (*, **, ***) indicate a p-value<0.05, <0.01, and <0.001 respectively, by Tukey’s HSD test.

We then assessed if GSDMD and MLKL-expressing cells were competent to induce autophagy when it is triggered by an external stimulus. Thus, we treated cells with rapamycin for 2 h after inducing the expression of the heterologous proteins for 5 h in galactose-containing media. Cells bearing an empty vector and treated for 2 h with rapamycin were used as a positive control of autophagy and untreated cells as a negative control. Neither cells expressing GSDMD(NT) nor cells expressing any of the MLKL constructs could induce autophagy with the same efficiency as control cells, although the effect was only statistically significant for cells expressing the different MLKL versions (Fig. 8B). We confirmed these results by visualizing Atg8-GFP localization in cells under the same conditions of rapamycin treatment (Fig. 8C). In control cells and cells expressing full-length GSDMD, GFP fluorescence mostly accumulated within the vacuole or in a single cluster per cell, consistent with the induction of autophagy. GSDMD(NT) did not seem to hamper accumulation of fluorescence in the vacuole as a readout of autophagy, while MLKL versions did lower the percentage of cells degrading Atg8-GFP. Yet, in cells expressing both GSDMD(NT) and all MLKL versions, we detected an increase in the percentage of cells that showed cytosolic Atg8 localization, revealing cells could not form autophagosomes at all in response to rapamycin. Moreover, indicating a problem in the traffic of autophagosomes towards the vacuole, MLKL constructs tended to accumulate multiple Atg8 clusters. This effect was significantly patent in the case of MLKL(1-182), as compared to GSDMD(NT). In conclusion, although to different degrees, GSDMD(NT) and MLKL disturb autophagic traffic, even though they should trigger autophagy as a consequence of TORC1 inhibition and Atg13 dephosphorylation. This could be explained if human pyropototic and necroptotic effectors caused a direct blockade of vesicular traffic.

### The NTDs of GSDMD and MLKL distinctly disrupt endosomal traffic

TORC1 is localized primarily on the vacuolar membrane and endosomes under basal conditions, where it interacts with EGOC to become activated. Previous reports highlight that homotypic fusion and protein sorting (HOPS), and class C core vacuole/endosome tethering (CORVET) complexes, as well as the endosomal sorting complex required for transport (ESCRT), all implicated in membrane and endosome fusion events, are important for the proper functioning of the TORC1 signaling pathway. Their disruption inhibits yeast growth even in the presence of nutrients [58–61]. The vesicular traffic machinery is also necessary to lead Atg8 to the vacuole during autophagy [62, 63]. Besides, several previous reports have related GSDMD and MLKL function to perturbations in the endosomal transport [20-22, 27-30]. Overall, our results suggested that GSDMD(NT) and all the MLKL constructs tested could be interfering with yeast endosomal traffic, consequently inhibiting TORC1 while hampering autophagy. To test this hypothesis, we studied whether endocytosis was altered by using the endocytic fluorescent marker FM4-64. GSDMD(NT) and the different constructs of MLKL interfered with proper traffic of FM4-64 to the vacuolar membrane (Fig. 9A-C), although some differences among them were noted. About 25% of GSDMD(NT)-expressing cells accumulated FM4-64 in the prevacuolar compartment (reminiscent of the well-established class E phenotype of yeast vacuolar protein sorting (*vps*) mutants), and another 25% displayed a diffuse FM4-64 signal throughout the cytoplasm (typical of a class C *vps* phenotype). In the case of MLKL, the three versions acquired a class C *vps*-like phenotype. The cause for each of these *vps* phenotypes is the alteration of distinct specific components of the endosomal pathway: the class C phenotype is associated with alterations in the HOPS complex [64, 65], involved in the fusion of late endosomes to the vacuoles, and the class E phenotype is associated with defects in the function of the ESCRT complexes [64, 66], involved in the formation of multivesicular bodies (MVBs).

**Figure 9.**
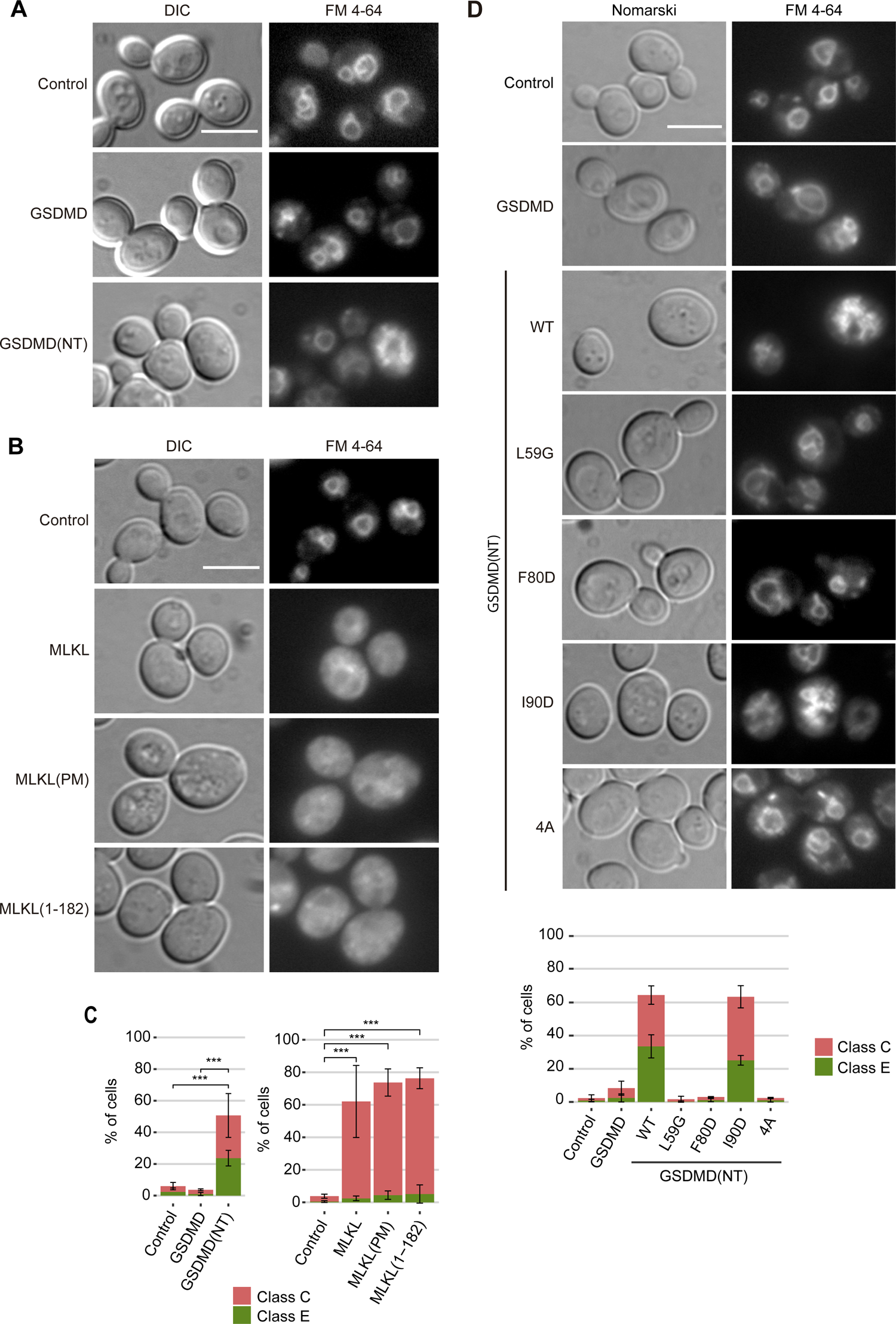
The NTDs of GSDMD and MLKL distinctly disrupt endosomal traffic. **(A-B)** Fluorescent and bright-field (DIC) microscopy of BY4741 strain bearing the same plasmids as in Fig. 1B and Fig. 1F, respectively, stained with FM4-64. **(C)** Quantification (n>100) of the percentage of cells showing a class C and E phenotype for each population shown in (A-B). **(D)** Fluorescent, bright-field (DIC) microscopy, and quantification (n>100) of the percentage of cells showing a class C and E *vps* phenotype for each population of BY4741 strain bearing the same plasmids as in Fig. 5A stained with FM4-64. Protein expression was induced for 5 h in SG medium in all cases. All scale bars indicate 5 µm. Results correspond to the mean of three biological replicates performed on different transformants. Error bars represent SD. Asterisks (***) indicate a p-value <0.001 by Tukey’s HSD test.

To verify these results, we assessed the impact of the expression of the GSDMD(NT) mutants described before on the endosomal traffic. As expected, like in the case of mitochondrial disruption, cells expressing the non-functional L59G, F80D, and 4A GSDMD(NT) mutants endocytosed the dye as efficiently as the control or full-length GSDMD, while cells expressing the functional GSDMD(NT) I90D behaved like GSDMD(NT) WT with a mixed class C and E *vps*-like phenotype (Fig. 9D).

Then, we evaluated vacuolar morphology using Vph1-GFP, a vacuolar membrane protein, as a marker. The results obtained supported our observations with FM4-64 (Fig. S7): in cells expressing GSDMD(NT), Vph1-GFP signal accumulated at the prevacuolar compartment in around 30% of the population, while cells expressing any of the versions of MLKL presented a mixed phenotype, with 10-15% of the cells displaying the same phenotype as GSDMD(NT) and a similar number showing disrupted vacuoles. Although we cannot discard completely that GSDMD(NT) or any of the constructs of MLKL are interacting directly with endosomal or vacuolar membrane granules, they did neither colocalize with endosomes nor with Vph1 (data not shown). These data support the hypothesis that TORC1 inhibition and the disruption of autophagy might be related to the interference of GSDMD(NT) and MLKL with endosomal traffic.

## Discussion

Here we report a yeast-based model for molecular studies on the effector proteins of pyroptosis and necroptosis. Despite the phylogenetic distance, heterologous expression in *S. cerevisiae* provides a ready experimental platform, a sort of an “*in vivo* cellular test tube” to elucidate particular mechanistic details on GSDMD and MLKL function. In the absence of homologous pathways, yeast supplies a cellular environment to study the fundamental properties of these pore-forming effectors. Furthermore, our previous work demonstrated that GSDMD can be processed by Caspase-1 in the yeast cell [39], and work by Ji *et al.*[40], proving that MLKL phosphorylation and activation can be recapitulated in yeast, opens the way for developing synthetic models to aid studies on these human cell death pathways.

To allow comparison of both human proteins, in the case of MLKL, we artificially produced the phosphomimetic mutant MLKL(PM), as well as a truncated MLKL(1-182) version, which should structurally compare to the constitutively active NTD of GSDMD released by Caspase-1 cleavage. Although it has been shown that human RIPK3 and MLKL co-expression in yeast enhances the intrinsic toxicity of the former in this model [40], in our hands a human phosphomimetic MLKL T357E/S358D [MLKL(PM)] behaved like the wild type. This agrees with studies arguing that mouse MLKL(PM) gains activity while the human protein mutated in the equivalent phosphorylation sites may not. Rather, in contrast to the mechanistic evidence raised in mice, it has been suggested that phosphorylation by RIPK3 in human cells may keep MLKL inactive [67]. This stresses the importance of developing alternative models to explore human proteins. Interestingly, the truncated MLKL(1-182) version was more active than full-length MLKL on yeast cells, even though MLKL does not undergo proteolysis for its activation in higher cells, unlike GSDMD. This stresses the idea that the C-terminal extension plays a regulatory role, as it does in GSDMD. The slight growth inhibition and cell death induced by MLKL and MLKL(PM) in yeast, compared to that of MLKL(1-182), may reflect either that the ability of the C-terminal pseudokinase extension of MLKL to block the interaction of its NTD with cellular membranes is less tight than in the case of GSDMD, or the existence of a phosphorylation-independent function for this protein [68]. Furthermore, contrary to what we observe for GSDMD, for which only the NTD alone forms aggregates detectable by immunoblot in non-reducing conditions, all MLKL different constructs aggregated, independently of their different ability to inhibit yeast growth. Some authors have claimed that MLKL exists as small oligomers under basal conditions that transit to high-order oligomers during necroptosis [69]. Our detection of aggregates may be consistent with this hypothesis.

These two pore-forming effector proteins mainly target the plasma membrane in higher eukaryotic cells, due to their capacity to interact with negatively charged phospholipids present in its inner layer, primarily phosphatidylserine and phosphoinositides such as phosphatidylinositol-4-phosphate and phosphatidylinositol-4,5-*bis*phosphate [4–6, 10]. In the yeast model, expression of the NTDs of both GSDMD and MLKL led to severe growth inhibition and loss of viability. However, we were unable to detect the association of EGFP fusions of both NTDs with the plasma membrane. Moreover, as loss of viability did not correlate with loss of plasma membrane selective permeability, we could not conclude that cell lysis was the main cause of toxicity. We cannot discard that the absence of severe membrane damage is a consequence of the presence of the bulky GFP tag, as previously described [47], because the fusion of GSDMD(NT) to a smaller FLAG tag significantly increased uptake of the vital marker propidium iodide. In any case, our data hint that the yeast model may be especially useful to study the interference of the pyroptotic and necroptotic pore-forming effectors with cytoplasmic membranes, as we found here that growth inhibition relates to interference with trafficking and TORC1 signaling.

As a proof-of-principle that the yeast model can be used to titrate the self-assembly of human GSDMD(NT) monomers, we developed and studied point mutants in residues equivalent to those described in the literature as implicated in interactions between monomers or with membrane phospholipids in murine GSDMD [10, 11]. Mutation of key residues in interface I (L59G), interface II (F80D), and the phospholipid-interacting region (4A) abolished GSDMD(NT) activity, while mutation of the interface III (I90D) had a milder impact. The reproduction of these interactions *in vivo* in yeast constitutes the first proof to our knowledge that such interfaces of contact are functionally conserved between the human and mouse proteins. Besides, the localization of loss-of-function GSDMD(NT) mutants at the mitochondria reveal that, even when they lose their capacity to aggregate, they can interact with particular lipid membranes, arguing in favor of a model in which the interaction with the membranes precedes oligomerization [3, 14, 70], or at least oligomerization is not a pre-requisite for membrane interaction. Surprisingly, the 4A mutant, lacking basic residues presumably involved in electrostatic interactions with membranes [10], also showed colocalization with mitochondria, although the signal was fainter than for the other mutants. At least two polybasic regions have been reported to be responsible for the interaction with phospholipids, which could explain this outcome when only one of them is mutated [11, 71]. GSDMD(NT) and MLKL(1-182) target yeast mitochondria, although we could only clearly colocalize MLKL(1-182) with this organelle. However, the fact that all loss-of-function GSDMD(NT) mutants colocalized with this structure in yeast strongly argues in favor of a direct association of GSDMD(NT) with mitochondrial membranes. The stronger fragmentation induced by WT GSDMD(NT) on yeast mitochondria as compared to MLKL(1-182) may be preventing the detection of this colocalization. Nevertheless, we cannot conclude that mitochondrial damage is a consequence of pore-forming activity. Previous studies have also reported a mitochondrial fragmentation effect in mammalian cells for both GSDMD(NT) and MLKL [6, 15–17, 19, 30]. The interaction between them and mitochondria is supported by the fact that they display a high affinity for cardiolipin [4–6, 10], although this lipid is present in the inner mitochondrial membrane and very scarce in the outer mitochondrial membrane [72]. Yeast cells show a slightly higher content of cardiolipin in mitochondria if compared to mammalian cells [73]. The first question that needs to be addressed is how these proteins reach the cardiolipin-rich membranes or which other lipids allow them to bind to the mitochondrial membrane. We tested the effects of cardiolipin removal in a *crd1*Δ yeast strain that lacks the cardiolipin synthase, the enzyme necessary for the synthesis of cardiolipin [72, 74], but we did not observe any changes in growth, localization, or mitochondrial damage (data not shown), so alternative mitochondrial outer membrane lipids might be involved.

Besides targeting mitochondria, both proteins seemed to impair endosomal traffic with some particularities: while GSDMD(NT) induced a mixed *vps-*type phenotype between class E and C (associated with a dysfunction of ESCRT and HOPS and complexes, respectively), MLKL induced a class C *vps* phenotype [64–66]. Our results suggest that each protein interferes with a particular point of the endosomal pathway by interacting with a protein or lipid present at that stage. Moreover, both proteins blocked autophagic flux, which is not surprising, as the ESCRT complex is involved in autophagosome closure and the HOPS complex in the delivery of their cargo into the vacuole [62, 63]. Endosomes are highly dynamic compartments [75], which can explain why we could not colocalize GSDMD or MLKL NTDs with endosomal or vacuolar membranes by microscopy. Like in the case of mitochondria, we cannot discard that GSDMD and MLKL interact directly with yeast endosomal and/or vacuolar membranes, perturbing them or even forming pore-like structures. It is noteworthy that all the MLKL versions cause the same damage in endosomal traffic while their phenotype on growth, cell death, and subcellular distribution are different. This implies that additive factors must be involved in the toxicity achieved by MLKL(1-182) (see below). Different studies have reported a relationship between GSDMD/MLKL and vesicular transport. In a screening aimed at identifying GSDMD(NT) regulators, several genes associated with the endosomal and vacuolar organization were detected in macrophages [21]. Another study identified that several proteins related to lysosomal function and trafficking were up-regulated in GSDMD-deleted osteoclasts [22]. Besides, the ESCRT-III system, which seems to be affected in yeast by GSDMD(NT) expression, is necessary for the repair of the plasma membrane and downregulation of pyroptosis [76]. As for MLKL, different authors have claimed that this protein might induce or inhibit autophagy, play a role in the formation of intraluminal vesicles in the MVB or induce its exocytosis, although further studies should confirm these results [20, 27–30, 77]. Besides, the ESCRT-III system plays a similar role during necroptosis to that described in the case of GSDMD-induced pyroptosis in the repair of the plasma membrane [78, 79]. Our results may help to clarify the specific vesicular traffic compartment that is targeted by these proteins.

MLKL and GSDMD(NT) show affinity for phosphoinositides, preferentially for phosphatidylinositol-4-phosphate [PI(4)P] and phosphatidylinositol-4,5-*bis*phosphate [PI(4,5)P_2_], typical of the plasma membrane [10, 80, 81], also for phosphatidylinositol-3-phosphate [PI(3)P] and phosphatidylinositol-3,5-*bis*phosphate [PI(3,5)P_2_], which are present in yeast early and late endosomes, respectively [5, 22]. As the concentration of PI(3)P and PI(3,5)P_2_ is higher in yeast endosomal compartments compared to those of mammalian cells [73], GSDMD and MLKL might be hijacked to endosomal vesicles in yeast, preventing them from localizing at the plasma membrane. Also, a blockade of trafficking caused by their presence could inhibit their own transport of pre-assembled aggregates to the plasma membrane by exocytosis. In any case, the differences in vesicle content and composition among the different types of mammalian cells might explain some of the controversies that have arisen on this subject.

An interesting finding is that the NTDs of pyroptotic and necroptotic effectors trigger TORC1 inhibition in yeast. Furthermore, our data suggest that loss of viability relies on cell cycle arrest as a consequence of TORC1 inhibition and the uncoupling of autophagy, rather than on cell lysis or organellar damage. Recent studies have established a link between GSDMD and pyroptosis to the mammalian TORC1 (mTORC1) pathway. Evavold *et al.* [21] showed that mTORC1 activity is necessary for the generation of ROS that drive GSDMD(NT) oligomerization. Other studies relate GSDMD activation by Caspase-8 to the Ragulator complex, the activator of mTORC1 [82, 83]. In our setting, our main hypothesis is that inhibition of TORC1 by both GSDMD(NT) and MLKL(1-182) is a consequence of the perturbation of the endosomal traffic rather than a direct interaction of GSDMD or MLKL with yeast TORC1 or any of its regulators. Vesicular traffic is involved in the interaction of TORC1 with both its activators and its substrates [58, 59, 61]. Recently, it was reported that there are two co-existing pools of TORC1 in yeast cells that regulate independent pathways. TORC1 located at the endosomes controls autophagy through Atg13 phosphorylation, while TORC1 located at the vacuolar membrane controls cell cycle progression through Sch9 phosphorylation [84]. Besides, a third pool of TORC1 is responsible for directly phosphorylating Maf1 within the nucleus to regulate RNA polymerase III, although it can also be phosphorylated by Sch9 [51, 53, 54]. Interestingly, although both GSDMD(NT) and MLKL(1-182) impaired cell cycle progression, they did not have the same impact on TORC1 activity: GSDMD(NT) caused a decrease in the phosphorylation of Sch9, Atg13, and Maf1, implying all TORC1 pools are affected; while MLKL(1-182) only affected Sch9 signaling, implying that only the TORC1 pool located at the vacuolar membrane is affected. It would be interesting to assess if these differences are associated with the distinct effect of GSDMD(NT) and MLKL(1-182) on endosomal traffic; and why MLKL and MLKL(PM), despite having a similar effect on endosomal traffic, did not affect TORC1 signaling significantly. It should not be overlooked that full-length MLKL, but not the NTD alone, induce proteostatic stress in the yeast cell, which may contribute for its distinct behavior. The tighter TORC1 inhibition caused by GSDMD(NT) might explain why this protein causes a more severe effect on cell growth and viability, whereas the more efficient inhibition of HOPS by MLKL may account for its more efficient impairment of autophagy. In any case, these results add evidence to the idea that GSDMD and MLKL might play roles in human cells beyond cell death, related to trafficking and response to nutrient or oxidative stress.

To summarize, we provide evidence that *S. cerevisiae* can be exploited as a model to study the effectors of pyroptosis and necroptosis, to deepen the molecular mechanisms of these proteins and the interaction between monomers. In our model, the NTDs of human GSDMD and MLKL are toxic to yeast cells because they form aggregates that affect mitochondria, endosomal traffic, autophagy, and cell cycle progression. Understanding its limitations, this model can be advantageous in future studies to identify new targets, perform structure-function assays, and eventually, test drugs that modulate their activity.

## Materials and methods

### Strains, media, and growth conditions

The BY4741 *S. cerevisiae* strain (*MATα his3Δ1 leu2Δ0 met15Δ0 ura3Δ0*) or its (BY4741 *trp1Δ::NatMX6* derivative (a gift from Á. Sellers-Moya, Complutense University of Madrid, Spain) [85] were used in all experiments unless otherwise stated. MVY04 strain (isogenic to BY4741, *VPH1-GFP-URA3*) was used to visualize the vacuolar membrane [39]. MVY07 strain (isogenic to BY4741, *HSP104-mCherry::KanMX*) was used to visualize Hsp104 and was obtained by amplifying *mCherry-KanMX* from the plasmid pAP17 (a gift from Jeremy Thorner, University of California, CA, USA), using primers Hsp104_mCh_Fw and Hsp104_mCh_Rv, and integrating the product in *HSP104* genomic locus. See Table 1 for primer sequences. The *Escherichia coli* DH5α strain was used for routine molecular biology techniques.

**Table 1.**
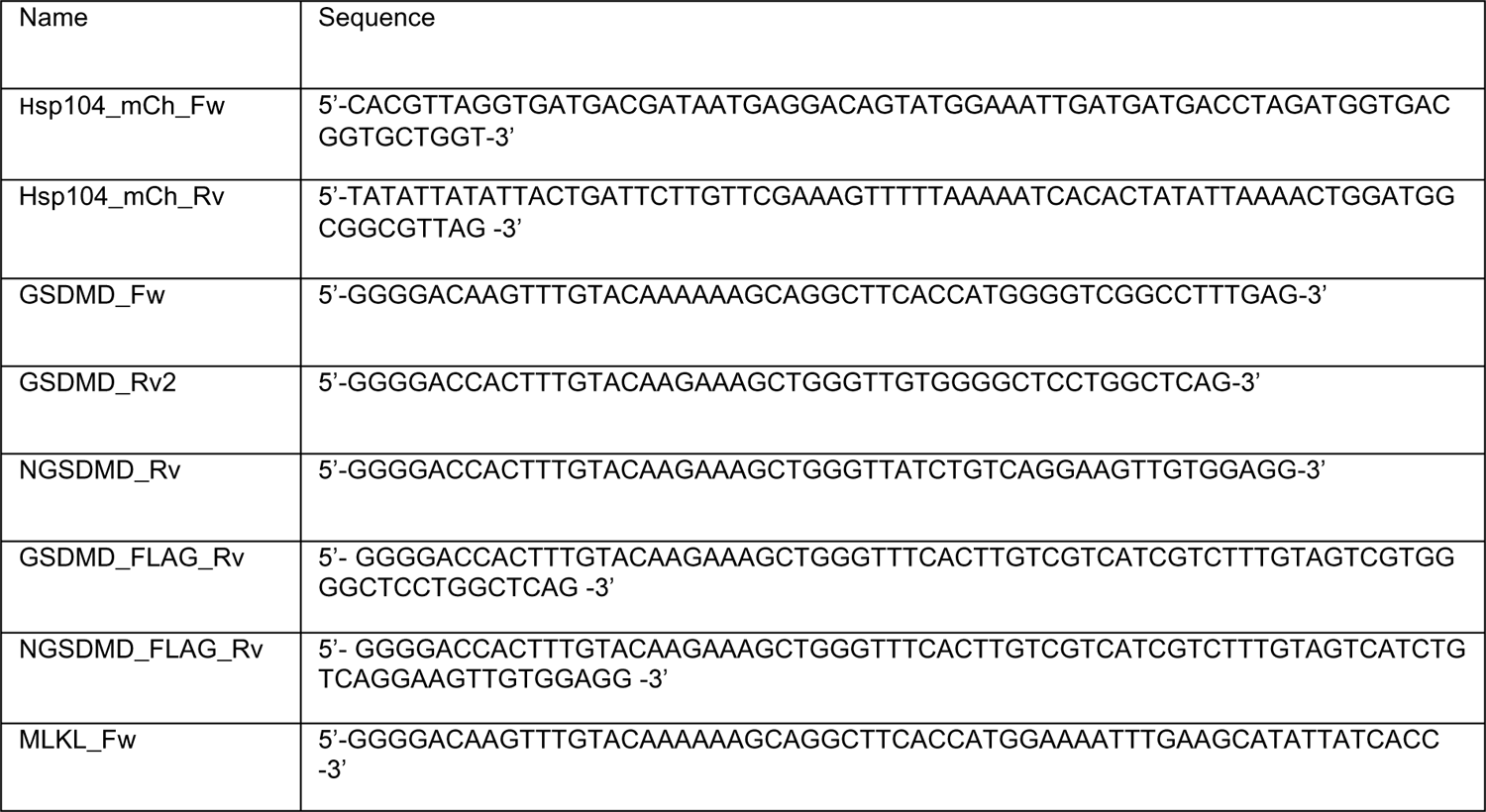
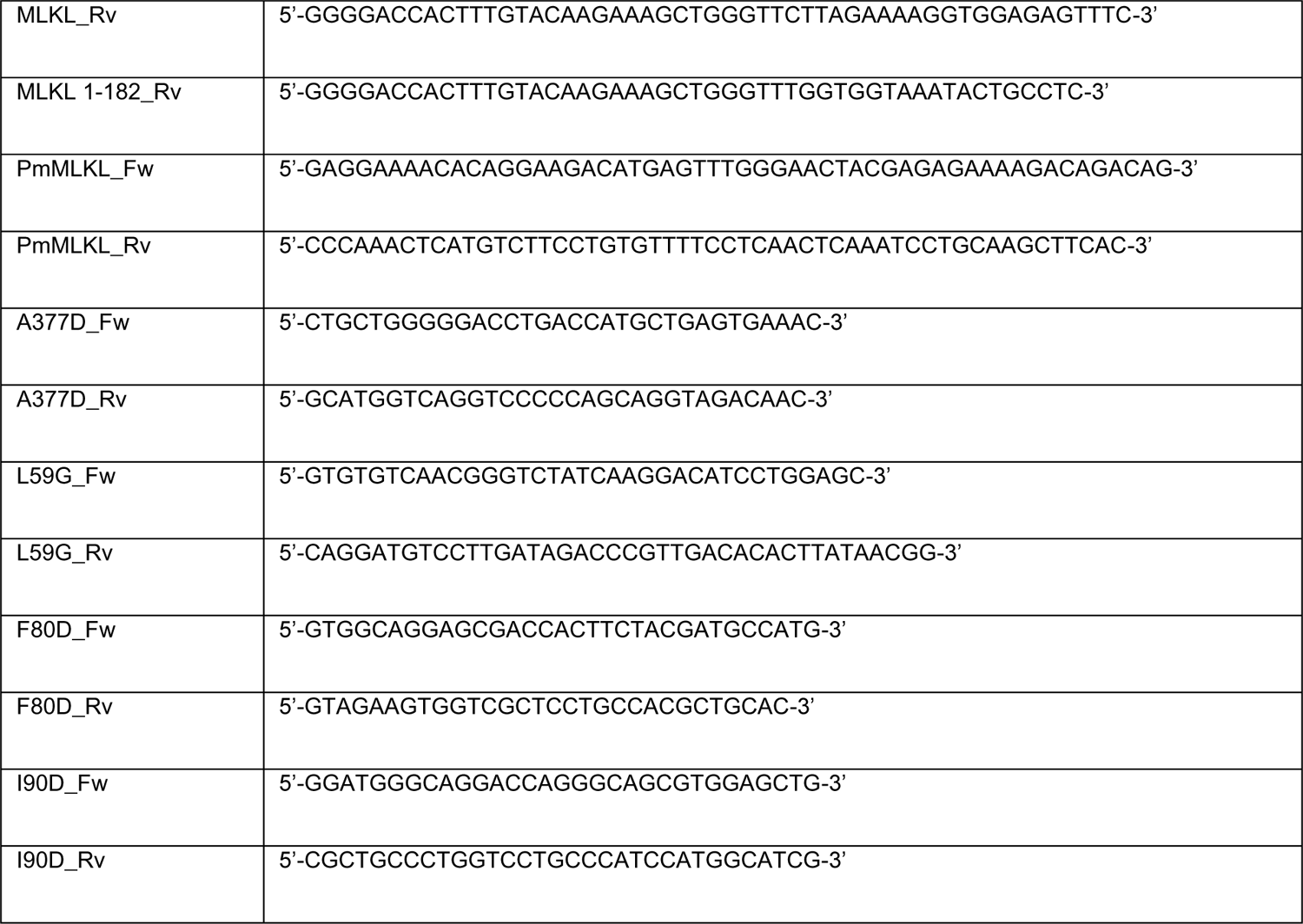
Oligonucleotides used in this work

Synthetic dextrose (SD) medium contained 2% glucose (ITW reagents), 0.17% yeast nitrogen base without amino acids (BD Difco), 0.5% ammonium sulfate (ITW reagents), and 0.12% synthetic amino acid drop-out mixture (Formedium), lacking appropriate amino acids and nucleic acid bases to maintain selection for plasmids. For synthetic galactose (SG) and synthetic raffinose (SR) media, glucose was replaced with 2% (w/v) galactose (ITW reagents) or 1.5% (w/v) raffinose (VWR), respectively. *GAL1*-driven protein induction in liquid medium was performed by growing cells in SR to mid-exponential phase and then refreshing the cultures to an OD600 of 0.3 directly with SG lacking the appropriate amino acids to maintain selection for plasmids for 5 h unless otherwise stated. Yeast strains were incubated at 30 °C.

### Plasmids

Transformation of *E. coli* and *S. cerevisiae* and other basic molecular biology methods were carried out using standard procedures.

*GSDMD* and *GSDMD(NT)* genes were amplified by standard PCR from pDB-His-MBP-GSDMD FL (a gift from J. Kagan, Boston Children’s Hospital, MA, USA) using primers GSDMD_Fw and GSDMD_Rv2 for the first, and GSDMD_Fw and NGSDMD_Rv for the second, all designed with *attB* flanking sites. *GSDMD-FLAG* and *GSDMD(NT)-FLAG* constructions were obtained by standard PCR from pDB-His-MBP-GSDMD FL using primers GSDMD_Fw and GSDMD_FLAG_Rv for the first, and GSDMD_Fw and NGSDMD_FLAG_Rv for the second, all designed with *attB* flanking sites. GSDMD R137A/K145A/R151A/R153A(NT) mutant, referred to as GSDMD(NT) 4A, was amplified by standard PCR from pDB-His-MBP-GSDMD 4A FL (a gift from J. Kagan, Boston Children’s Hospital, MA, USA) using the same primers as for the wild-type gene. *MLKL* gene and its truncated version *MLKL(1-182)* were amplified by standard PCR from pRetrX-TRE3G-hMLKL-Venus (Addgene_106078) using primers MLKL_Fw and MLKL_Rv for the first, and MLKL_Fw and MLKL 1-182_Rv for the second. See Table 1 for primer sequences. The *attB*-flanked PCR products were cloned into pDONR221 vector by Gateway BP Clonase II reaction (Invitrogen) to generate entry clones. Subsequently, the inserts from the entry clones were subcloned into pAG413-GAL-ccdB-EGFP/DsRed, pAG415-GAL-ccdB, or pAG416-GAL-ccdB-EGFP vectors (Addgene_1000000011) [86] by Gateway LR Clonase II reaction (Invitrogen), generating the plasmids pAG413-GSDMD-EGFP/DsRed, pAG413-GSDMD(NT)-EGFP/DsRed, pAG413-MLKL-EGFP/DsRed, and pAG413-MLKL(1-182)-EGFP/DsRed, pAG415-GSDMD-FLAG, pAG415-GSDMD(NT)-FLAG, pGA415-GSDMD(NT) 4A-FLAG pAG416-GSDMD-EGFP, pAG416-GSDMD(NT)-EGFP, pAG416-GSDMD(NT) 4A-EGFP, pAG416-MLKL-EGFP, and pAG416-MLKL(1-182)-EGFP. All the proteins were tagged in C-terminal, with a 17-amino acid linker (MVSKGEELFTGVVPILV) due to the characteristics of Gateway Cloning system. Only in -FLAG constructs the tag was fused immediately after the protein.

MLKL(PM), GSDMD A377D, and GSDMD(NT) L59G, I80D or I90D mutants were obtained by site-directed mutagenesis performed on their respective wild-type entry clone, using primers PmMLKL_Fw, PmMLKL_Rv, A377D_Fw, A377D_Rv, L59G_Fw, L59G_Rv, F80D_Fw, F80D_Rv, I90D_Fw, and I90D_Rv, respectively. Primers are listed in Table 1. Subsequently, the inserts from the entry clone were subcloned into pAG413-GAL-ccdB-EGFP/DsRed, pAG415-GAL-ccdB, and pAG416-GAL-ccdB-EGFP plasmids by Gateway LR Clonase II reaction, generating the plasmids pAG413-MLKL(PM)-EGFP/DsRed, pAG415-GSDMD(NT) L59G, F80D, or I90D-FLAG, pAG416-MLKL(PM)-EGFP, pAG416-GSDMD A377D-EGFP, and pAG416-GSDMD(NT) L59G, F80D, or I90D-EGFP. As stated above, all the proteins were tagged at their C-terminal ends.

pJU676 (pRS416-Sch9-5xHA) and pAH099 (pRS416-MAF1-3xHA) plasmids, used as a readout for TORC1 activity, were a gift from R. Loewith, University of Geneva, Switzerland [50]. HC078 (pRS315-3xHA-Atg13) plasmid, used also as a readout of TORC1 activity, was obtained from Addgene (Addgene_ 59544). The autophagic marker Atg8-GFP, encoded in the plasmid pRS314-GFP-Atg8, was a gift from Y. Ohsumi, Tokyo Institute of Technology, Japan [87]. The mitochondrial marker Ilv6-mCherry, encoded in the plasmid pOB06 was a gift from Ó. A. Barbero, Complutense University of Madrid, Spain.

### Western blotting assays

Western blotting assays were carried out by standard techniques in 10% acrylamide (ITW reagents) gels [39]. Non-reducing western blots were performed by removing dithiothreitol (DTT) from the sample buffer and using 7.5% acrylamide gels. Assessment of Sch9 phosphorylation was adapted from Péli-Gulli *et al.* [88]. Twenty mL of cell culture were mixed with trichloroacetic acid (TCA) (Sigma-Aldrich) at a final concentration of 6% and incubated in ice for 10 min. After centrifugation, the pellet was washed with ice-cold acetone and dried in a SpeedVac SC100 (Savant). The pellet was resuspended in a volume of urea buffer [50 mM Tris-HCl pH 7.5 (Fisher BioReagents), 6 M urea (Merck), 1% sodium dodecyl sulfate (SDS) (Duchefa Biochemie), 50mM NaF (Probus), and 1mM phenylmethanesulfonyl fluoride (PMSF) (Amresco) proportional to the OD600nm of the cell culture. Cells were disrupted by bead beating with FastPrep24 (MP Biomedicals). Subsequently, 2X sample buffer was added [120 mM Tris-HCl pH 6.8, 20% glycerol (ITW Reagents), 200 mM DTT (Acros), 4% SDS] and the mix was boiled at 60°C for 10 minutes. Proteins were resolved by SDS-PAGE in 7.5% acrylamide gels and transferred onto nitrocellulose membranes for 1 h 30 min at 80 V.

Mouse anti-GFP (BD Biosciences JL-8,1:1,000 dilution) and anti-HA (Sigma-Aldrich 12CA5, 1:1,000 dilution) were used as primary antibodies to detect the expression of proteins fused to GFP and HA, respectively. Rabbit anti-glucose-6-phosphate dehydrogenase (G6PDH) antibody (Sigma-Aldrich, 1:50,000 dilution) was used as a loading control. Anti-rabbit IgG-IRDye 800CW (LI-COR Biosciences), anti-rabbit IgG-IRDye 680LT (LI-COR Biosciences), anti-mouse IgG-IRDye 800CW (LI-COR Biosciences), anti-mouse IG-IRDye 680LT (LI-COR Biosciences), all at 1:5,000 dilution, were used as secondary antibodies. Odyssey XF Imaging System (LI-COR Biosciences) or ChemiDoc MP Imaging System (Bio-Rad) were used for developing the immunoblots.

### Spot growth assays

Spot growth assays on plates were performed by incubating transformant clones overnight in SR media, adjusting the culture to an OD_600_ of 0.5, and spotting samples in four serial 10-fold dilutions onto the surface of SD or SG plates lacking the appropriate amino acids to maintain selection for plasmids, followed by incubation at 30 °C for 2-3 days.

### Microscopy techniques

For *in vivo* bright field differential interference contrast (DIC) microscopy or fluorescence microscopy, cells were cultured as previously stated, harvested by centrifugation at 3,000 rpm for 3 min, and viewed directly on the microscope. Cells were examined with an Eclipse TE2000U microscope (Nikon) using the appropriate sets of filters. Digital images were acquired with an Orca-ER camera controller (Hamamatsu) and were processed with the HCImage software.

Observation of actin in yeast cells with Rd-conjugated phalloidin (Invitrogen) was performed as previously described [39]. For FM4-64 (Invitrogen) vital staining, cells were cultured as previously stated, harvested by centrifugation, and resuspended in synthetic medium. Cells were labeled with 2.4 µM FM4-64, incubated for 1 h 15 min at 30 °C with shaking, washed in FACSFlow™ Sheath Fluid (BD Biosciences), and observed by fluorescence microscopy.

For confocal microscopy, cells were cultured as previously stated, harvested by centrifugation, and fixed with a 4% p-formaldehyde (ITW Reagents), and 3.4% sucrose (ITW Reagents) solution for 15 min at room temperature. Then cells were washed and resuspended in FACSFlow™ Sheath Fluid. Coverslips were treated with 5 µL of concanavalin A (Sigma-Aldrich) and dried at room temperature. Adhesion of cells was performed by adding 7 µL of fixed cells over concanavalin A-treated coverslips and incubating for 30 min. ProLong^TM^ Glass Antifade Mountant (ThermoFisher Scientific)/Glycerol (1:1) was used to avoid photobleaching. Cells were examined with an Olympus IX83 Automated Fluorescence Microscope, coupled to Olympus FV1200 confocal system, using the appropriate set of filters. Images were processed to remove background and enhance contrast. Images were analyzed using Fiji and Adobe Photoshop.

### Flow cytometry

Cells were cultured as previously stated. After 5h of galactose induction, cell death measurement by PI (Sigma-Aldrich) staining was performed as previously described [39]. For cell cycle analysis, 1.5×10^7^ cells were harvested, fixed, and permeabilized with 70% ethanol overnight for each sample. Then samples were treated at 37 °C for 2 h with 500 µL of RNAse A (Roche) 2 mg/mL and after 30 min with 200 µL of pepsin (Sigma-Aldrich) 10 mg/mL. DNA was stained by the addition of 0.0005% PI in FACSFlow™ Sheath Fluid.

Cells were analyzed using a FACScan (Becton Dickinson) flow cytometer through a 585/42 BP emission filter (FL2) for PI. At least 10,000 cells were analyzed for each experiment. Data were processed using FlowJo software.

### Cell viability assay

Cells were cultured as previously stated. After 5 h of galactose induction, cell viability was measured by the microcolonies method, as described before [39, 89].

### Statistical analysis

Data were analyzed using RStudio, ggplot2, dplyr, tidyverse, ggrepel, openxlsx, ggthemes, ggsignif, gridExtra, and Origin software. All data sets were tested for normality using the Shapiro-Wilkinson test. When a normal distribution was confirmed, a One-Way ANOVA test with a *post hoc* Tukey’s HSD test was used for statistical comparison between multiple groups. For data sets that did not show normality, a Kruskal-Wallis test with a *post hoc* Dunn’s test was applied. The asterisks (*, **, ***) in the figures correspond to a p-value of <0.05, <0.01, and <0.001, respectively. Experiments were performed as biological triplicates on different clones and data with error bars are represented as mean ± standard deviation (SD).

### Structure analysis

The schemes of GSDMD and MLKL structure were generated using Illustrator for Biological Sequences (IBS) [90]. The alignment of GSDMD sequences was done using the Clustal Omega (EMBL-EBI). The 3D projections of GSDM3A were built using PyMOL.

## Acknowledgments

We thank Á. Sellers-Moya, Ó. A. Barbero, R. Loewith, J. Thorner, and Y. Ohsumi for materials; and J. Kagan for materials and useful discussion; C. Mazzoni, C. Evavold, and our colleagues at Research Unit 3 for their support and discussion; and L. Sastre for technical support. M. V. was supported by a predoctoral contract from Universidad Complutense de Madrid. We thank the Genomics Unit (Genomics and Proteomics Center, UCM) for their help with the sequencing reactions, the Confocal and Multiphoton Microscopy Unit (Cytometry and Fluorescence Microscopy Center, UCM) for their help with the confocal microscopy experiments, and the Flow Cytometry Unit (Cytometry and Fluorescence Microscopy Center, UCM) for their help with the flow cytometry experiments. This research was possible thanks to funding from Grant PID2019-105342GB-I00 from Ministerio de Ciencia e Innovación (Spain) to M. M. and V. J. C.

## Author contribution

M. V.: Conceptualization, Methodology, Validation, Formal analysis, Investigation, Writing – original draft, and Visualization. M. M. and V. J. C.: Conceptualization, Methodology, Resources, Writing – Review & Editing, Supervision, Project administration, and Funding acquisition.

