## Supplementary Figures for "Human Gasdermin D and MLKL disrupt mitochondria, endocytic traffic and TORC1 signaling in budding yeast"

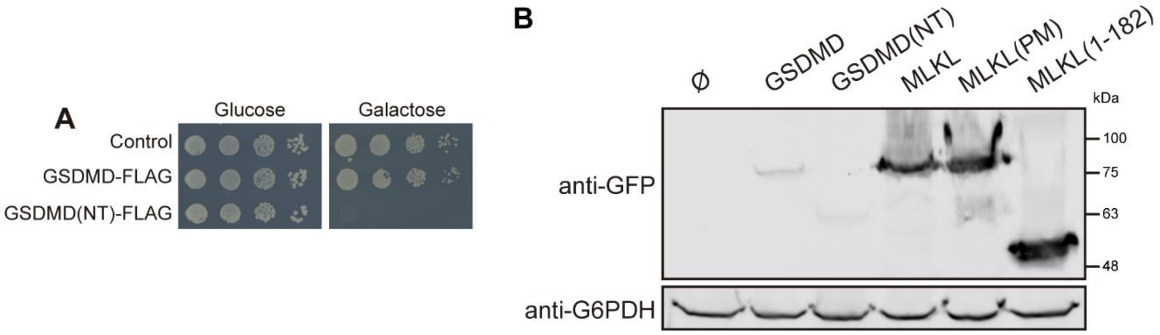


**Figure S1.** The NTD of GSDMD fused to a FLAG tag inhibits yeast growth, related to Fig. 1. **(A)** Spot growth assay of BY4741 strain bearing plasmids pAG415-GSDMD-FLAG and pAG415-GSDMD(NT)-FLAG. pAG415 empty vector was used as a control. Cells were cultured on SD (Glucose) and SG (Galactose) agar media for repression and induction of GSDMD and GSDMD(NT) expression, respectively. **(B)** GSDMD is expressed in yeast in much lower levels than MLKL. Immunoblot showing a comparison of the levels of expression of the different constructs of GSDMD and MLKL using yeast lysates of BY4741 strain bearing plasmids pAG416-GSDMD-EGFP, pAG416-GSDMD(NT)-EGFP, pAG416-MLKL-EGFP, pAG416-MLKL(PM)-EGFP, and pAG416-MLKL(1-182)-EGFP after 5 h induction in SG medium. pAG416-EGFP empty vector was used as a control. The membrane was hybridized with anti-GFP antibody. Anti-G6PDH antibody was used as a loading control. A representative assay from three different experiments with different transformants is shown in all cases.

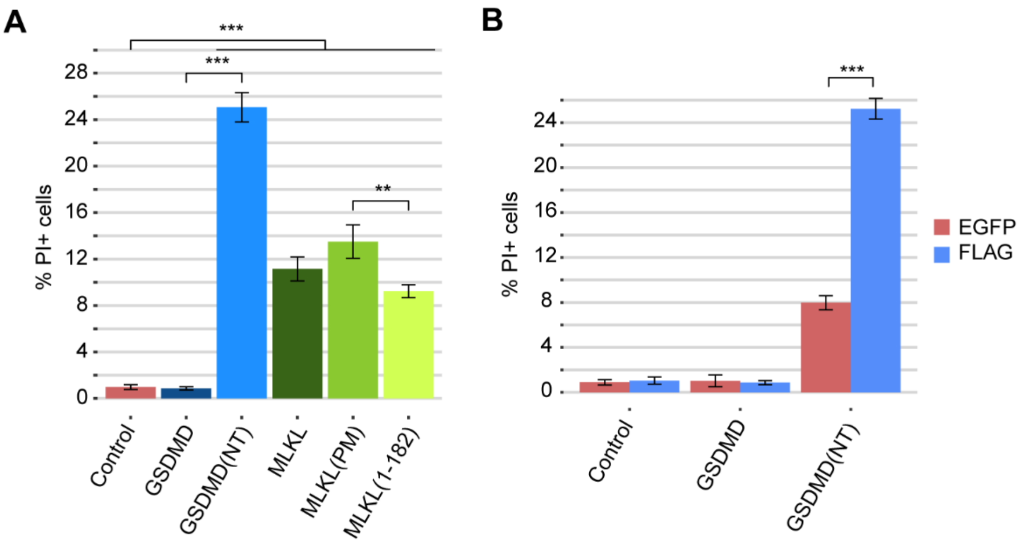


**Figure S2.** The NTDs of GSDMD and MLKL induce cell death, related to Fig. 3. **(A)** Graph showing the percentage of PI-positive stained cells (n=10,000) for each population of BY4741 strain bearing plasmids pAG416-GSDMD-EGFP, pAG416-GSDMD(NT)-EGFP, pAG416-MLKL-EGFP, pAG416-MLKL(PM)-EGFP, and pAG416-MLKL(1-182)-EGFP after 12 h of induction in SG medium. pAG416-EGFP empty vector was used as a control. **(C)** Graph showing the percentage of PI-positive stained cells (n=10,000) for each population of BY4741 strain bearing plasmids pAG415-GSDMD-FLAG, pAG416-GSDMD-EGFP, pAG415-GSDMD(NT)FLAG, and pAG416-GSDMD(NT)-EGFP after 5 h of induction in SG medium. pAG415 and pAG416-EGFP empty vectors were used as controls. Results correspond to the mean of three biological replicates performed on different transformants in all cases. Error bars represent SD. Asterisks (**, ***) indicate a p-value <0.01 and <0.001, respectively, by Tukey's HSD test.

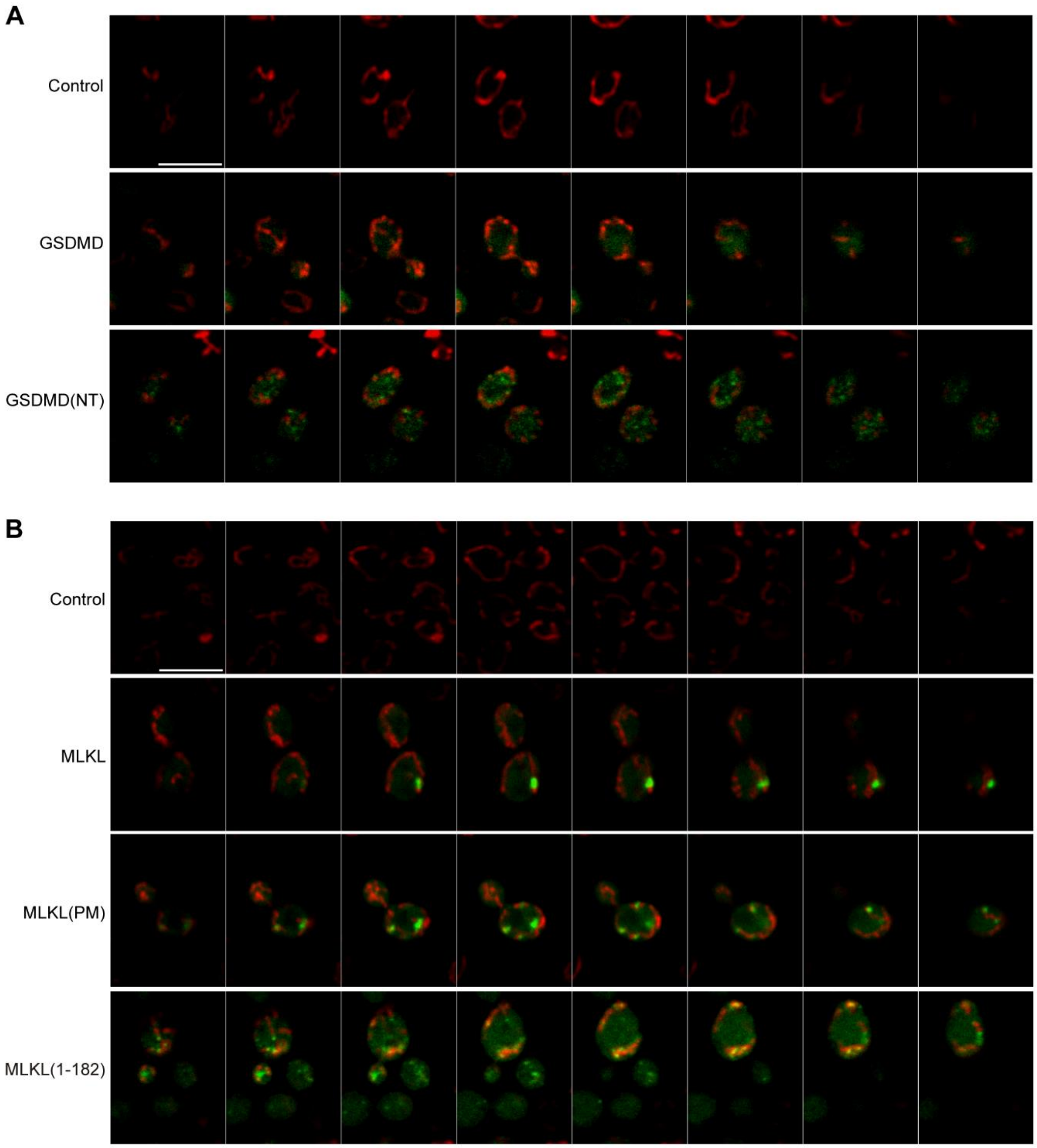


**Figure S3.** The NTDs of GSDMD and MLKL fragment the mitochondrial network, related to Fig. 4. **(A)** Confocal merged slices of BY4741 strain bearing the mitochondrial marker pOB06 (Ilv6-mCherry) and plasmids pAG416-GSDMD-EGFP and pAG416-GSDMD(NT)-EGFP, respectively. pAG416 empty vector was used as a control. **(B)** Confocal merged slices of BY4741 strain bearing the mitochondrial marker pOB06 (Ilv6-mCherry) and plasmids pAG416-MLKL-EGFP, pAG416-MLKL(PM)-EGFP and pAG416-MLKL(1-182)-EGFP, respectively. pAG416 empty vector was used as a control. All scale bars indicate 5 μm. Protein expression was induced for 5 h in SG medium in all cases

**A**

```
mGSDMA3(NT)  -MPVFEDVTRALVRELN-PRGDLTPLDSLIDFKHFRPFCLVLRKRKSTLFWGARYVRTDY  58
 hGSDMD(NT)  MGSAFERVVRRVVQELD-HGGEFIPVTSLQSSTGFQPYCLVVRKPSSSWFWKPRYKCVNL  59
 mGSDMD(NT)  MPSAFEKVVKNVIKEVSGSRGDLIPVDSLRNSTSFRPYCLLNRKFSSSRFWKPRYSCVNL  60
                .** *.: :::*:.   *:: *: **  . . *:*:**: ** .*: **  **  .:

mGSDMA3(NT)  TLLDLLEPGSSPSDLTDSGNFSFKNMLDVQVQGLVEVPKT--VKVKGTAGLSQS--STLE  114
 hGSDMD(NT)  SIKDILEPDAAEPDVQRGRSFHFYDAMDGQIQGSVELAAPGQAKIAGGAAVSDSSSTSMN  119
 mGSDMD(NT)  SIKDILEPSAPEPEPECFGSFKVSDVVDGNIQGRVMLSGMGEGKISGGAAVSDSSSASMN  120
             :: *:***.:   :     .* . : :* ::** * :       *: * *.:*:*  ::::

mGSDMA3(NT)  VQTLSVAPSALENLKKERKLS-ADHSFLNEMRYHEKNLYVVMEAVEAKQEVTVEQTG--N  171
 hGSDMD(NT)  VYSLSVDPNTWQTLLHERHLRQPEHKVLQQLRSRGDNVYVVTEVLQTQKEVEVTRTHKRE  179
 mGSDMD(NT)  VCILRVTQKTWETMQHERHLQQPENKILQQLRSRGDDLFVVTEVLQTKEEVQITEVHSQE  180
             *  * *  .: :.: :**:*   :...*:::* : .:::** *.:::::** : ..   :

mGSDMA3(NT)  ANAIFSLPSLALLG--LQGSLNNNKAVTIPKGCVLAYRVRLLRVFLFNLWDIPYICNDSM  229
 hGSDMD(NT)  GSGRFSLPGATCLQGEGQGHLSQKKTVTIPSGSTLAFRVAQL--VIDSDLDVLLFPDKKQ  237
 mGSDMD(NT)  GSGQFTLPGALCLKGEGKGHQSRKKMVTIPAGSILAFRVAQL--LIGSKWDILLVSDEKQ  238
             ... *:**.   *    :*  ..:* **** *. **:**  *  .: .  *:  . :..

mGSDMA3(NT)  QTFPKIRRVPCSAFISPTQMISEEPEEEKLI------G-------- 261
 hGSDMD(NT)  RTFQPPATGHKRS--TSEGAWPQLPSGLSMMRCLHN------FLTD 275
 mGSDMD(NT)  RTFEPSSGDRKAV--GQ------RHHGLNVLAALCSIGKQLSLLSD 276
             :**                        .::
```

**B**

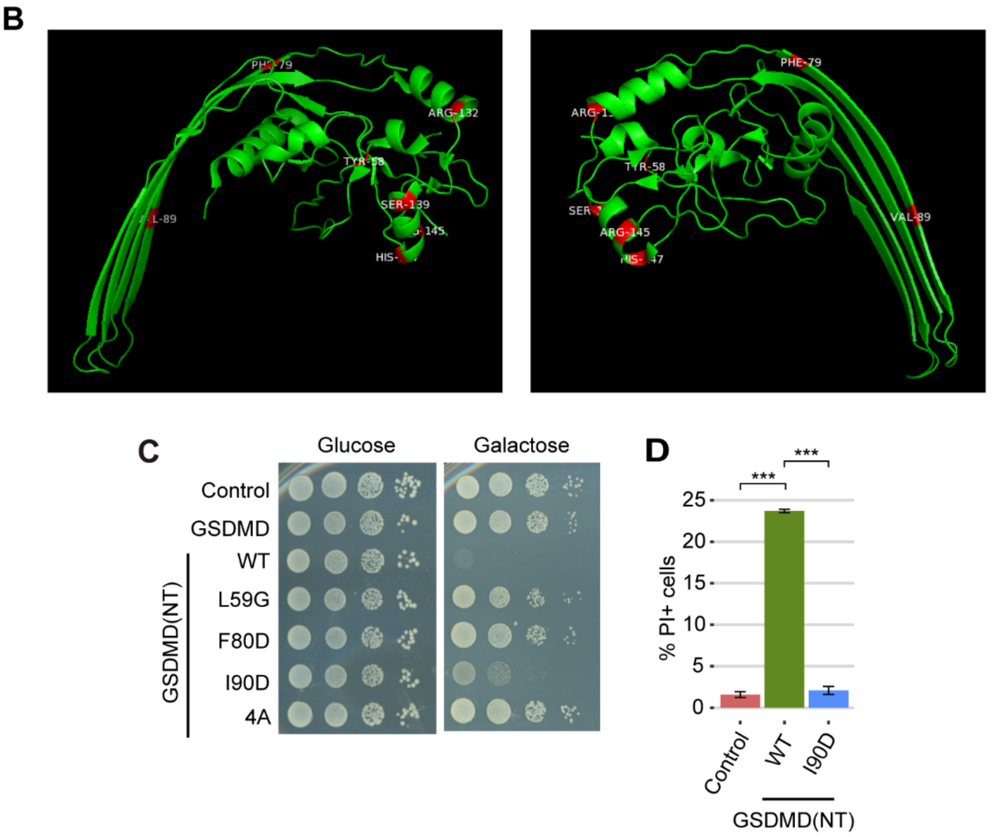


**Figure S4.** Structure-function analysis of GSDMD(NT), related to Fig. 5. **(A)** Alignment of the sequences corresponding to the NTD of mouse gasdermin A3 (mGSDMA3) (UPI000044F464), human GSDMD (hGSDMD) (UPI00001291ED), and mouse GSDMD (mGSDMD) (UPI0000021F53). The residues L50G, F80D, and I90D of hGSDMD(NT) that belong to the interface I, II, and III of interaction between GSDMD(NT) monomers, respectively, are highlighted in red. The residues R137, K145, R151, and R153 of hGSDMD(NT) involved in the interaction of GSDMD(NT) with membrane phospholipids are highlighted in green. **(B)** 3D projections of the hGSDMD residues indicated in (A) on the structure of mGSDMA3 NTD (ePDB 6cb8). **(C)** Spot growth assay of BY4741 strain bearing plasmids pAG415-GSDMD (NT) WT, F80D, I90D, or 4A-FLAG. pAG415 empty vector was used as a control. Cells were cultured on SD (Glucose) and SG (Galactose) agar media for repression and induction of GSDMD and GSDMD(NT) expression, respectively. **(D)** Graph showing the percentage of PI-positive stained cells (n=10,000) for each population of BY4741 strain bearing the same plasmids as in (C) after 5 h of induction in SG media. Results correspond to the mean of three biological replicates performed on different transformants in all cases. Error bars represent SD. Asterisks (***) indicate a p-value <0.001 by Tukey's HSD test.

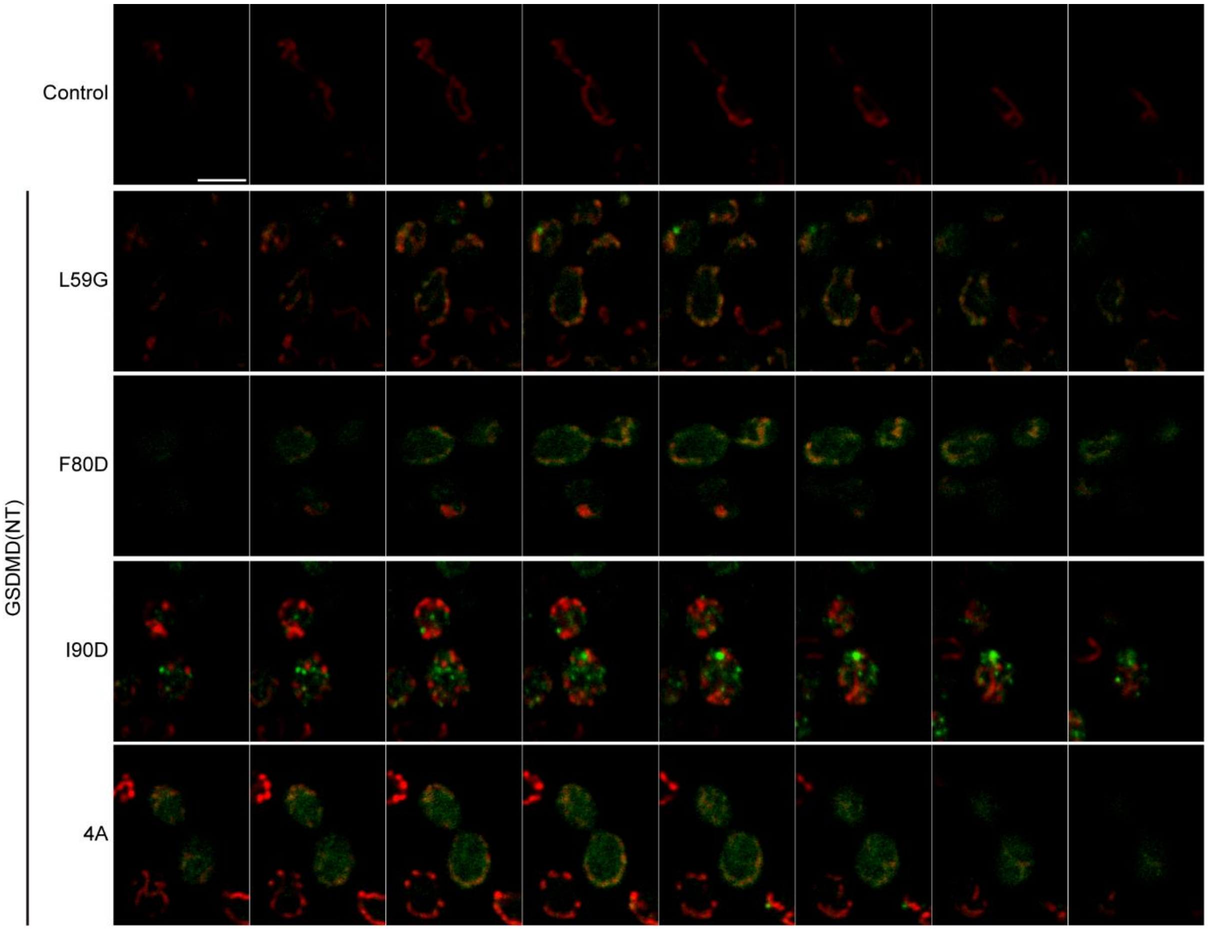


**Figure S5.** Non-toxic mutants of the NTD of GSDMD colocalize with the mitochondrial network, related to Fig. 6**.** Confocal merged slices of BY4741 strain bearing the mitochondrial marker pOB06 (Ilv6-mCherry) and plasmids pAG416-GSDMD(NT)-EGFP L59G, F80D, I90D or 4A, after 5h of induction in SG medium. pAG416-Ø empty vector was used as a control. Scale bar indicates 5 µm.

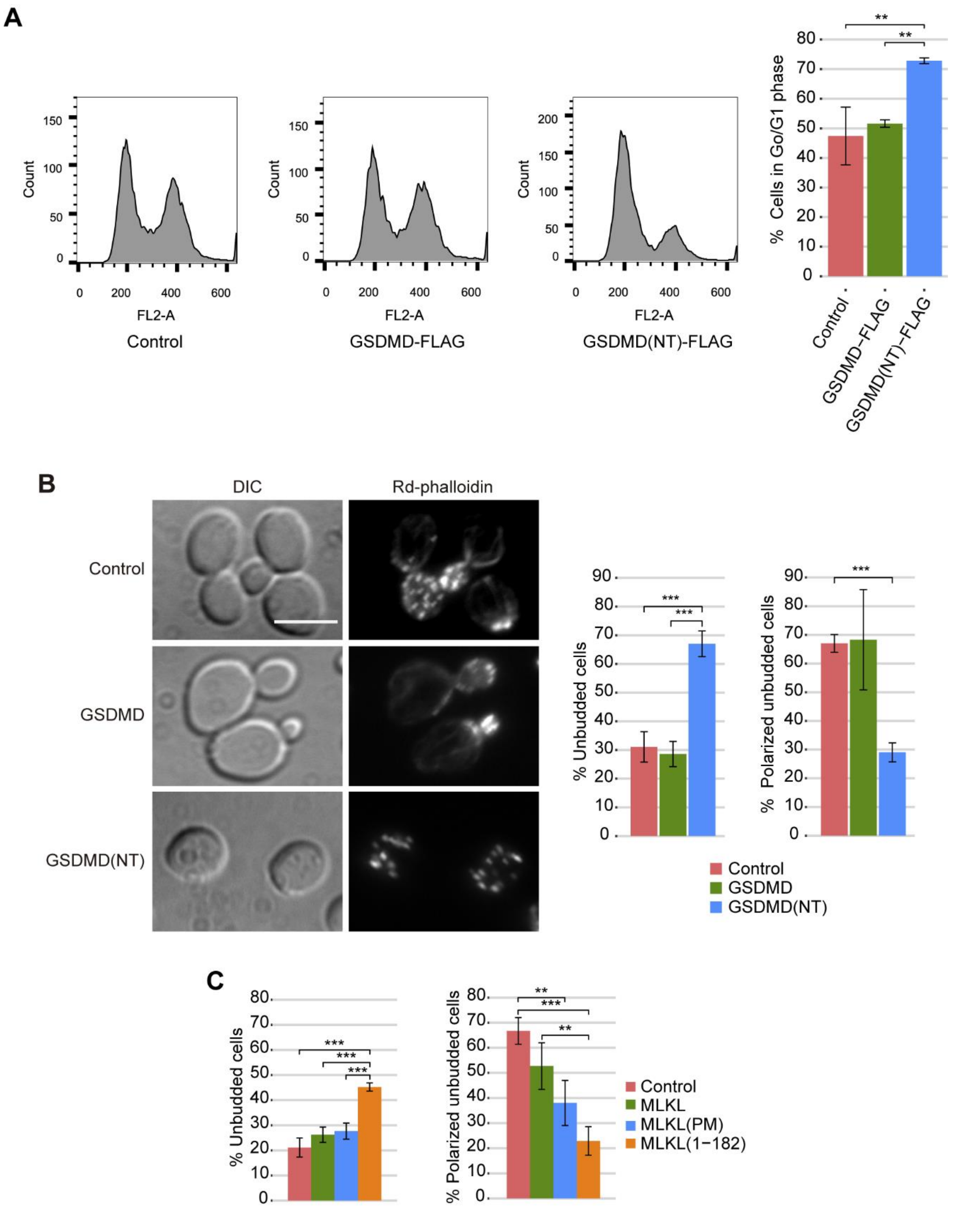


**Figure S6.** The NTDs of GSDMD and MLKL cause a cell cycle arrest, related to Fig. 7. **(A)** Cell cycle profiles obtained by measuring DNA content (FL2-A) of cells stained with PI and subsequently analyzed by flow cytometry (n=10,000) (left panels), and graph showing the percentage of cells in phase G0/G1 for each population (right panel) of BY4741 strain bearing plasmids pAG415-GSDMD-FLAG and pAG415-GSDMD(NT)-FLAG. pAG415 empty plasmid was used as a control. **(B)** Actin staining with Rd-phalloidin, bright field (DIC) microscopy, and quantification (n > 100) of the percentage of unbudded cells and the percentage of cells with polarized cytoskeleton among the unbudded cells of BY4741 strain bearing pAG416-GSDMD-EGFP and pAG416-GSDMD(NT)-EGFP after 5 h of induction in SG medium. pAG416-EGFP empty vector was used as a control. Scale bar indicates 5 µm. **(C)** Quantification (n > 100) after actin staining of the percentage of unbudded cells and the percentage of cells with polarized cytoskeleton among the unbudded cells of BY4741 strain bearing the plasmids pAG416-MLKL-EGFP, pAG416-MLKL(PM)-EGFP, and pAG416-MLKL(1-182)-EGFP, after 5 h of induction in SG medium. pAG416-EGFP empty vector was used as a control. Protein expression was induced for 5 h in SG medium in all cases. Results correspond to the mean of three biological replicates performed on different transformants. Error bars represent SD. Asterisks (**, ***) indicate a p-value <0.01 and <0.001, respectively, by Tukey's HSD test.

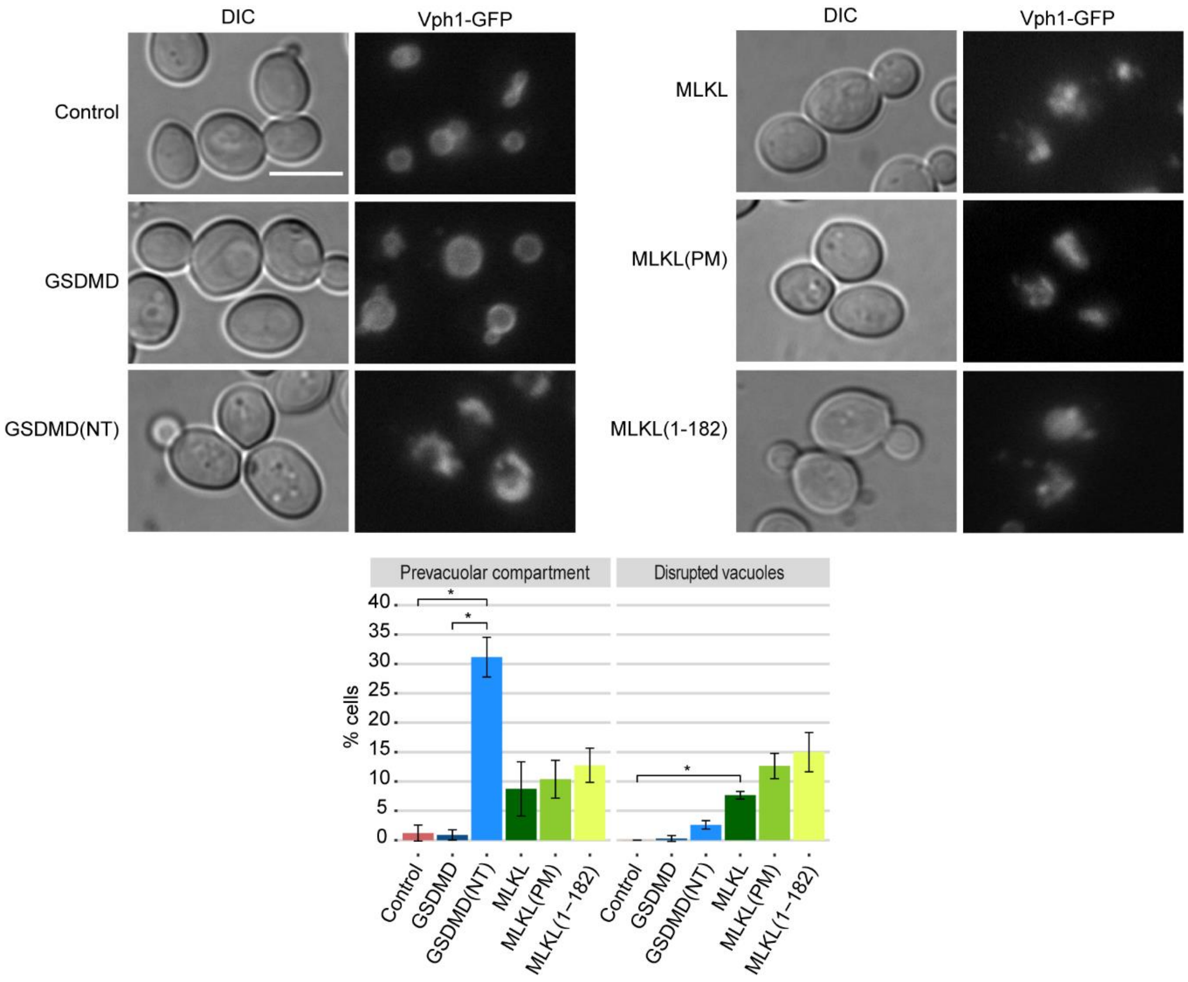


**Figure S7.** GSDMD(NT) and the different constructs of MLKL interfere with the yeast vacuole, related to Fig. 9. Fluorescent and bright-field (DIC) microscopy (upper panel) and quantification (n > 100) (lower panels) of the vacuolar phenotype of MVY04 strain bearing the plasmids pAG413-GSDMD-DsRed, pAG413-GSDMD(NT)-DsRed, pAG413-MLKL-DsRed, pAG413-MLKL(PM)-DsRed or pAG413-MLKL(1-182)-DsRed after 5 h of induction in SG medium. pAG413-DsRed empty vector was used as a control. Scale bar indicates 5 µm. Results correspond to the mean of three biological replicates performed on different transformants. Error bars represent SD. Asterisks (*) indicate a p-value <0.05 by Tukey's HSD test.
